# Stereospecific GPG acylation by CLN8 drives BMP biosynthesis and its loss leads to Batten disease

**DOI:** 10.64898/2025.12.18.695077

**Authors:** Pradeep K. Sheokand, Denis Lacabanne, Andrew M. James, Stefania Della Vecchia, Jonathan J. Ruprecht, Joris van der Kleij, Keira Turner, Jessica Müller-Niva, Miia H. Salo, Benjamin Jenkins, Susannah K. Leese, Nidhi Juneja, Chak Shun Yu, Clarissa D. Booth, Martin S. King, Johanna Uusimaa, Jill M. Weimer, Albert Koulman, Reetta Hinttala, Filippo M. Santorelli, Maria Marchese, Michael P. Murphy, Edmund R. S. Kunji, Kasparas Petkevicius

## Abstract

Loss-of-function mutations in the endoplasmic reticulum membrane protein CLN8 cause Batten disease, a neurodegenerative lysosomal storage disorder^1^. Together with the lysosomal enzyme CLN5, CLN8 mediates the biosynthesis of bis(monoacylglycero)phosphate (BMP), a phospholipid essential for lysosomal function and distinguished by its unique *S,S* stereochemistry^2,3^. However, the role of CLN8 in BMP synthesis has remained unclear. Here we establish that CLN8 is a glycerophosphoglycerol (GPG) acyltransferase that catalyses the stereospecific acylation of *S,S*-GPG to produce *S,S*-lysophosphatidylglycerol (LPG), the CLN5 substrate in BMP synthesis. Using cryo-electron microscopy, we resolve structures of the CLN8 homodimer in apo and substrate-bound states at 2.7 Å resolution, revealing the active site architecture and a ping-pong acyl transfer mechanism. Batten disease-causing missense mutations impair CLN8 enzymatic activity in vitro and reduce BMP levels in a Cln8*^R24G^* knock-in mouse, whereas the Cln8*^mnd^* mouse frameshift mutation causes complete loss of BMP in vivo. Exogenous *S,S*-LPG, but not the *R,S* stereoisomer, restored BMP synthesis in CLN8- deficient cells and mice, and improved neurological phenotypes in *cln8* mutant zebrafish. Together, these findings define the enzymatic function of CLN8, elucidate the biochemical basis of CLN8 Batten disease, and establish a proof-of-concept for treating it through stereospecific BMP precursor supplementation.

## Main

Batten disease, or neuronal ceroid lipofuscinosis, encompasses a group of inherited lysosomal storage disorders caused by mutations in 13 distinct *CLN* genes^1^. Despite this diversity of genetic causes, most forms share a hallmark pathology: impaired lysosomal function leading to the accumulation of undegraded lipids and proteins as lipofuscin^4^. This buildup drives neuroinflammation and neurodegeneration, with symptoms typically emerging in late infancy and progressing to premature death early in life^4^. The precise molecular roles of many CLN proteins in lysosomal homeostasis, however, are poorly defined.

Bis(monoacylglycero)phosphate (BMP) is a phospholipid found exclusively in late endosomes and lysosomes, where it plays a key role in maintaining lysosomal membrane dynamics and catabolic capacity^5^. Unlike other animal phospholipids, which contain a glycerol backbone with an *R*-configured chiral centre, BMP consists of two glycerol molecules linked to a phosphate group, each containing a chiral carbon in the *S* configuration (*S,S*-BMP)^6–8^. Cellular BMP is derived from phosphatidylglycerol (PG), which features an *R*-configured backbone and an *S*-configured glycerol headgroup (*R,S*-PG)^9–13^. The pathway converting *R,S*-PG into *S,S*-BMP has yet to be defined.

Two Batten disease proteins, CLN5 and CLN8, have recently been implicated in distinct acyl transfer steps within BMP biosynthesis^2,3^. CLN5 functions as a lysosomal transacylase, transferring an acyl group from one lysophosphatidylglycerol (LPG) molecule to the headgroup of another to complete the final step of BMP synthesis^3^. We previously identified CLN8, a member of the TRAM-LAG1-CLN8 (TLCD) acyltransferase family, as essential for cellular BMP production^2^. However, the precise acyl transfer reaction catalysed by CLN8, its molecular mechanism, and its position within the BMP biosynthetic pathway have remained unknown.

### CLN8 acylates *S,S*-glycerophosphoglycerol

To define the enzymatic role of CLN8 in BMP biosynthesis, we investigated its function in the context of CLN5. CLN5 acts as a lysosomal transacylase, and CLN5-deficient cells accumulate lysosomal LPG while lacking *S,S*-BMP^3,14^. We reasoned that because transacylation does not alter the glycerol backbone, the CLN5 substrate LPG must already have the *S,S* stereoconfiguration, placing CLN8 upstream in the conversion of *R,S*-PG to *S,S*-LPG (Fig. 1a).

**Fig. 1:**
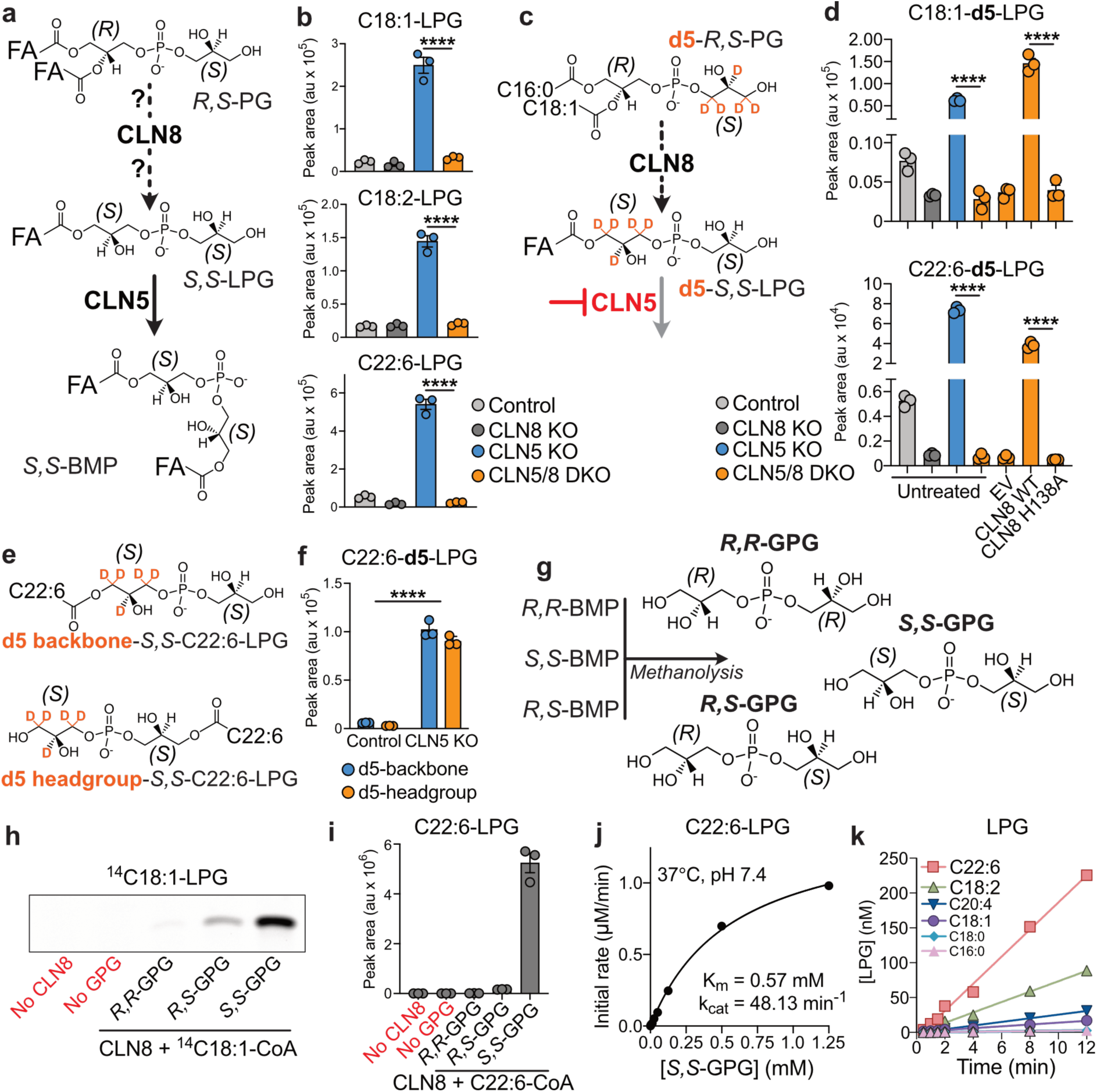
CLN8 is a stereospecific *S,S*-GPG acyltransferase in the BMP synthesis pathway. a, BMP biosynthesis pathway schematic. *R,S*-PG is converted to *S,S*-LPG through several steps, one of which is catalysed by CLN8. *S,S*-LPG then serves as both acyl donor and acceptor for CLN5 to generate *S,S*-BMP. **b,** The levels of the indicated LPG species in control, CLN8 KO, CLN5 KO, and CLN5/8 DKO HeLa cells under standard culture conditions (n = 3 per genotype). **c,** Schematic of the labelling experiment to trace the synthesis of *S,S*-LPG. **d,** Levels of d5-labelled LPG species in the indicated cell lines, either untreated or transfected for 48 h with CLN8 WT or the catalytically inactive H138A mutant, then treated with 25 μM d5-PG for 6h (n = 3). **e,** Structures of *S,S*-C22:6-LPG species containing the d5 label either in the glycerol headgroup or backbone. **f,** Abundance of C22:6-LPG species containing d5 label in the headgroup or backbone in control and CLN5 KO cells following 6 h labelling with 25 μM d5-PG (n = 3 per genotype). **g,** Schematic of the generation of *R,R*-, *R,S*-, and *S,S*-GPG substrates. **h,** Phosphorimaging of TLC-separated LPG band from in vitro assays with purified CLN8, ¹⁴C18:1-CoA, and different GPG stereoisomers. **i,** Abundance of C22:6- LPG product from in vitro assays using purified CLN8, C22:6-CoA, and different GPG stereoisomers. **j,** Michaelis-Menten kinetics of purified CLN8 activity using saturating C22:6- CoA and increasing concentrations of *S,S*-GPG. **k,** Time-dependent production of indicated LPG species in reactions with purified CLN8, equimolar acyl-CoA mix, and *S,S*-GPG. In vitro assay results are representative of at least two independent protein preparations. ****P < 0.0001 by one-way ANOVA with Dunnett’s post hoc test (**b, d**), or by two-way ANOVA (**f**).

To test this hypothesis, we utilised our previously published CLN8 knockout (KO) HeLa cells, which lack BMP species^2^, and generated CLN5/8 double knockout (DKO) cells by introducing a CLN5 deletion; in parallel, we engineered CLN5 single-KO cells (Extended Data Fig. 1a-c).

Comparison of LPG levels across these lines showed that CLN5 KO cells accumulated multiple LPG species, consistent with a block in the final pathway step converting LPG to BMP (Fig. 1b). This accumulation was lost in CLN5/8 DKO cells, demonstrating that CLN8 is required for LPG synthesis upstream of CLN5 (Fig. 1b).

To trace the conversion of *R,S*-PG to *S,S*-LPG directly, we supplemented cells with *R,rac-*PG containing a deuterium-labelled *racemic* glycerol headgroup and an *R*-configured diacylglycerol (C16:0/C18:1) backbone (d5-PG) (Fig. 1c). In CLN5 KO cells, d5-LPG species containing oleate (C18:1) or docosahexaenoate (DHA, C22:6) accumulated substantially, as was observed for unlabelled LPG species (Fig. 1b,d). This accumulation was lost in CLN5/8 DKO cells and restored by hemagglutinin (HA)-tagged wild-type CLN8 (CLN8-HA), but not by the catalytically dead CLN8 H138A mutant lacking the conserved TLCD histidine required for acyl transfer (Fig. 1d; Extended Data Fig. 1d)^2^. Thus, CLN8 acyltransferase activity is essential for *R,S*-PG to LPG conversion in cells.

To define the CLN8 reaction further, we analysed C22:6-d5-LPG, which accumulates in CLN5 KO cells but is depleted in CLN8 KO and CLN5/8 DKO cells (Fig. 1d). This lipid provides a specific readout of cellular CLN8 activity, as it can only arise through complete removal of the original 16:0 and 18:1 acyl chains from d5-PG, followed by CLN8-mediated C22:6-CoA acylation. Supplementation with d5-PG could, in principle, yield two forms of C22:6-d5-LPG: backbone-labelled and headgroup-labelled (Fig. 1e), which are distinguishable by their daughter ions in tandem MS (Methods). Surprisingly, tandem MS analysis revealed equal accumulation of both backbone- and headgroup-labelled C22:6-d5-LPG in CLN5 KO cells (Fig. 1f). This finding supports only a model in which CLN8 acylates a symmetrical substrate with two equivalent glycerol moieties – namely, glycerophosphoglycerol (GPG).

Although acylation of glycerophosphodiesters (GPDs) has previously been described only in yeast^15,16^, and GPDs in vertebrates are thought to be destined for degradation^17^, our data suggested GPG is a *bona fide* CLN8 substrate. To test this, we generated all three potential GPG stereoisomers (*S,S*; *R,S*; *R,R*) by deacylating synthetic BMP standards and validated them by LC-MS/MS against a chemically synthesised *R,S*-GPG standard (Fig. 1g). In in vitro assays with purified CLN8 (Extended Data Fig. 2a,b) and ¹⁴C18:1-CoA, CLN8 acylated *S,S*-GPG most efficiently, with substantially lower activity for *R,S*-GPG and only trace activity with *R,R*- GPG (Fig. 1h). Similar stereospecificity was observed using unlabelled C22:6-CoA, with robust C22:6-*S,S*-LPG production only from *S,S*-GPG (Fig. 1i). CLN8 acylated *S,S*-GPG with C18:1-CoA and C22:6-CoA in a time- and concentration-dependent manner, with substantially higher capacity than what we had previously observed for *R,racemic*-LPG acyl acceptor substrate (Fig. 1j; Extended Data Fig. 2c,d)^2^. Kinetic analysis showed CLN8’s affinity and catalytic efficiency for *S,S*-GPG was comparable to those of other acyltransferases acting on physiological substrates (Fig. 1j)^2,15^. CLN8 also displayed a marked preference for polyunsaturated acyl-CoA substrates, particularly C22:6-CoA and C18:2-CoA (Fig. 1k).

Lysosomal BMPs contain fatty acids esterified to the *sn*-2 and *sn’*-2 positions of the glycerol moieties (Extended Data Fig. 2e)^18^. This configuration implies that CLN8 should generate *S,S*- 2-acyl-LPG intermediates, positioning the acyl chain correctly for CLN5 to condense them into 2,2ʹ-BMP (Extended Data Fig. 2e). To determine whether CLN8 acylates the *sn*-2 or *sn*-3 position of the glycerol moiety in *S,S*-GPG, we analysed 1-C18:1- and 2-C18:1-LPG standards, which are readily resolved by LC-MS/MS (Extended Data Fig. 2f). When *S,S*-GPG was incubated with purified CLN8 and C18:1-CoA, the reaction product co-eluted with 2-C18:1- LPG standard (Extended Data Fig. 2g). Consistently, the C18:1-LPG species that accumulates in CLN5-deficient cells was also identified as 2-C18:1-LPG (Extended Data Fig. 2h). Together, our stable-isotope tracing and enzymatic assays identify CLN8 as a stereospecific GPG acyltransferase that converts *S,S*-GPG and acyl-CoAs into *S,S*-2-acyl-LPG. This product is then utilised by CLN5 in the lysosome to form *S,S*-2,2ʹ-BMP.

### Structure and reaction mechanism of CLN8

To dissect the biochemical mechanism of CLN8, we purified tag-cleaved, codon-optimised human CLN8 from yeast for structural studies (Extended Data Fig. 3a,b). Like other TLCD family members, purified CLN8 formed SDS-resistant multimers on SDS-PAGE, but size-exclusion chromatography (SEC) revealed a single homogeneous population (Extended Data Fig. 3c,d)^2^. The SEC eluate was concentrated, confirmed to be enzymatically active, and used for cryo-EM. Two-dimensional class averages revealed a dimeric architecture resembling its paralog ceramide synthase 6 (CERS6) (Extended Data Fig. 4a)^19^. The initial structure, resolved at 3.65 Å (Supplementary Table 1), showed well-defined transmembrane domains, whereas the ER-luminal N-terminus (residues 1–15) and cytosolic C-terminus (residues 261–286) were disordered and not resolved (Extended Data Fig. 4a, Extended Data Fig. 5a)^20^. CLN8 forms a homodimer, with each monomer containing seven transmembrane helices and an interface mediated by TM6 (Fig. 2a). Unexpectedly, we observed two coenzyme A (CoA) molecules bound in pockets located within the transmembrane region of each monomer, positioned at the dimer interface (Fig. 2a; Extended Data Fig. 5a). Given its polarity, CoA incorporation within the membrane likely occurs during dimer assembly.

**Fig. 2:**
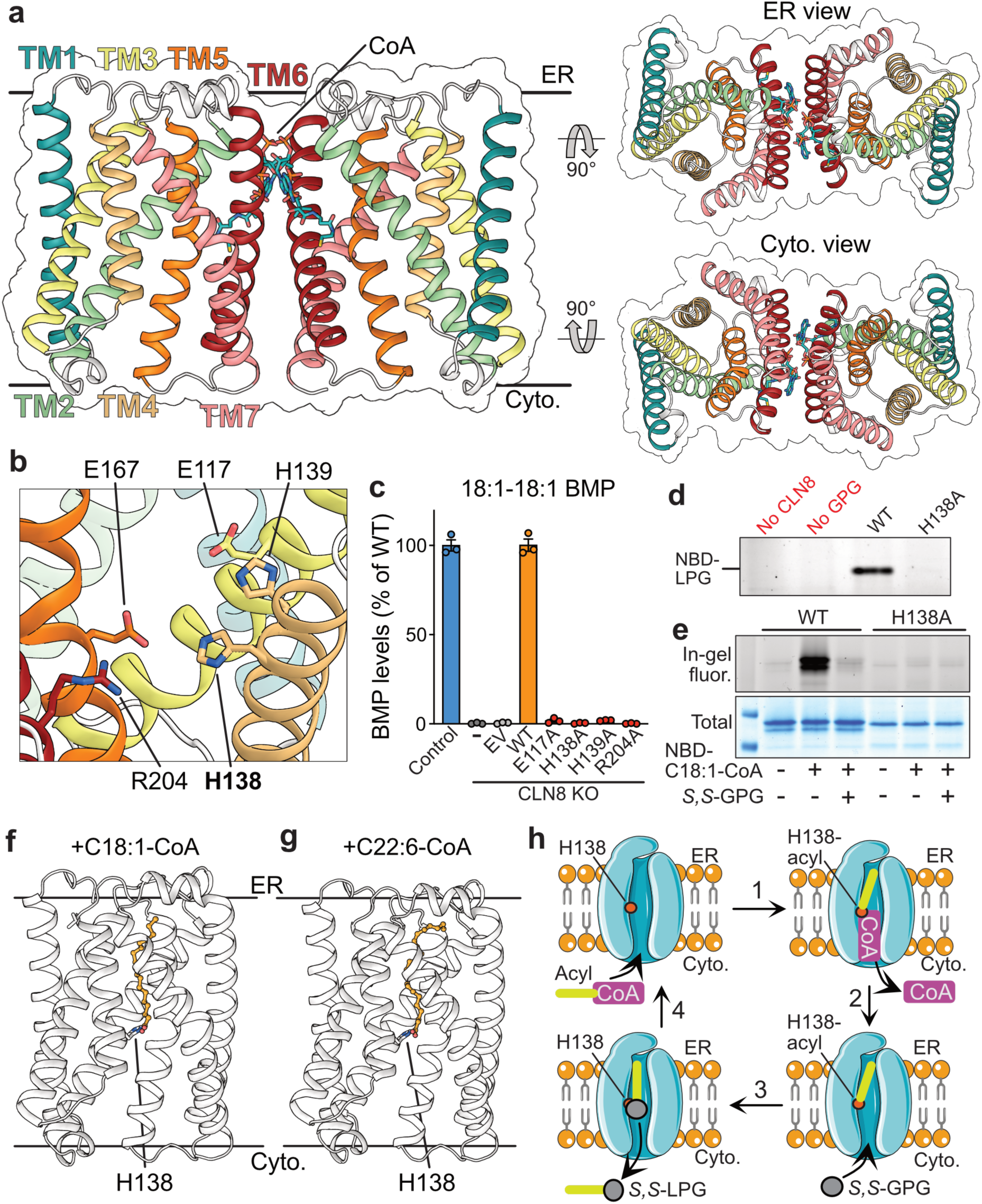
Structure and reaction mechanism of CLN8 homodimer. a, Cartoon representation of the CLN8 homodimer, shown in side view and in views from the ER lumen and cytoplasm, with bound CoA molecules in cyan. **b,** Active site of CLN8 highlighting catalytic residues. **c,** Cellular levels of 18:1-18:1 BMP in control and CLN8-knockout HeLa cells transfected with CLN8 WT or alanine mutants of active-site residues (n=3). **d,** Fluorescence scan of thin-layer chromatography showing LPG formation in in vitro assays with NBD-18:1-CoA, purified CLN8 WT or H138A, and *S,S*-GPG. **e,** In-gel NBD fluorescence (top) and Coomassie staining (bottom) of SDS–PAGE of purified CLN8 WT and H138A incubated with NBD–18:1–CoA alone or together with S,S-GPG, as indicated. **f–g,** CLN8 cartoon structures solved with C18:1–CoA and C22:6–CoA, showing bound fatty acyl groups and H138 residues. Only one monomer of the dimer is shown. **h,** Catalytic cycle of CLN8, shown for clarity using only one monomer of the dimer. In vitro assay results are representative of at least two independent protein preparations.

The TLC domain (TM2–7) forms a barrel enclosing a central cavity open to the cytosol and sealed toward the ER lumen by the TM2–TM3 and TM4-TM5 loops (Fig. 2a). At the centre of this cavity, we identified a putative active site comprising E117, H138, H139, E167, and R204, which is conserved across the TLCD acyltransferases (Fig. 2b, Extended Data Fig. 1d)^2^.

Alanine substitutions of these residues (except E167A, which was not expressed) failed to restore BMP levels in CLN8 KO HeLa cells despite normal protein expression levels (Fig. 2c, Extended Data Fig. 1e), confirming their essentiality for CLN8 catalysis.

Given that the conserved histidines H211 in CERS6 and H117 in TLCD1 form covalent acyl-enzyme intermediates during ceramide and phosphatidylethanolamine synthesis^2,19^, respectively, we tested whether the analogous residue H138 in CLN8 serves the same role (Extended Data Fig. 1d). Wild-type CLN8, but not H138A, purified from human cells could utilise fluorescent NBD-C18:1-CoA as an acyl donor to catalyse NBD-C18:1-*S,S*-LPG synthesis from *S,S*-GPG (Fig. 2d). Consistent with formation of a catalytic intermediate, SDS-PAGE revealed a covalent fluorescent NBD-C18:1-CLN8 adduct that was lost upon addition of *S,S*- GPG, whereas no such intermediate was detected for H138A (Fig. 2e).

To further visualise this intermediate state, we determined cryo-EM structures of CLN8 prepared in the presence of either C18:1-CoA or C22:6-CoA at 2.7 Å resolution (Fig. 2f,g; Extended Data Fig. 4b,c; Supplementary Table 1). The improved resolution relative to the apo structure likely reflects stabilisation by the bound acyl chain, enabling refinement of the model (Fig. 2a; Extended Data Fig. 4; Extended Data Fig. 5b,c). In both complexes, the acyl chain was positioned with a carboxy group adjacent to H138 within the central cavity and the tail oriented toward the ER lumen (Fig. 2f,g). CoA molecules were also observed between the two monomers in the same configuration as in the apo structure (Extended Data Fig. 5b,c). In the C18:1-bound complex, we observed a single acyl chain density, whereas in the C22:6-bound complex, additional weaker densities suggested alternative binding poses for the flexible polyunsaturated chain (Extended Data Fig. 5d). The absence of CoA-SH in the active site suggests that these structures capture an acyl-enzyme intermediate poised to accept *S,S*-GPG from the cytosolic side (Fig. 2f,g). Although continuous density between H138 and the acyl chain was not resolved (Extended Data Fig. 5b,c), a covalent H138-acyl linkage is thermodynamically required for catalysis after CoA-SH dissociation from the active site. This is consistent with the formation of NBD-C18:1-CLN8 adduct and with homology to CERS6 and TLCD1^2,19^. Further supporting this model, when we added *S,S*-GPG to CLN8 that had been incubated with C22:6-CoA, we could only observe the protein in the apo state in cryo-EM, consistent with turnover and *S,S*-LPG product release. Together, these observations support a ping-pong acyl-transfer mechanism in which CLN8 catalysis proceeds via a covalent H138–acyl intermediate, analogous to other TLCD acyltransferases (Fig. 2h).

### Functional impact of Batten disease-causing CLN8 mutations

Next, we examined whether Batten disease-causing CLN8 missense mutations impair its enzymatic activity. We selected 11 mutations spanning a range of disease onset and severity, diverse amino acid chemistries, and distributed across different regions of the protein (Fig. 3a)^21–29^. Mutations affecting conserved active-site residues that we showed above are essential for catalysis (H139Y, R204C, and R204L; Fig. 2b,c) were not analysed^30–32^. Using plasmid-based overexpression, we first assessed whether these mutant variants altered CLN8 expression or the ability to restore BMP levels in CLN8 KO HeLa cells. Given the strong plasmid-driven overexpression, which produces protein levels far exceeding endogenous CLN8, the assay functioned as a binary test of residual activity: even substantially less active variants would be expected to restore BMP to wild-type levels. Two variants, A30P and L207R, were not expressed, likely due to protein destabilisation (Extended Data Fig. 6a). Of the remaining mutants, T170R and G237R failed to restore BMP in CLN8 KO cells, indicating a complete loss of activity (Extended Data Fig. 6b). Both uncharged residues are positioned adjacent to the active site, where substitution with a bulky, positively charged arginine likely perturbs the local electrostatic environment and thereby abolishes catalysis (Fig. 3a; Extended Data Fig. 6c). The other variants restored BMP in CLN8 KO cells, suggesting partial activity, and were therefore selected for further in vitro characterisation.

**Fig. 3:**
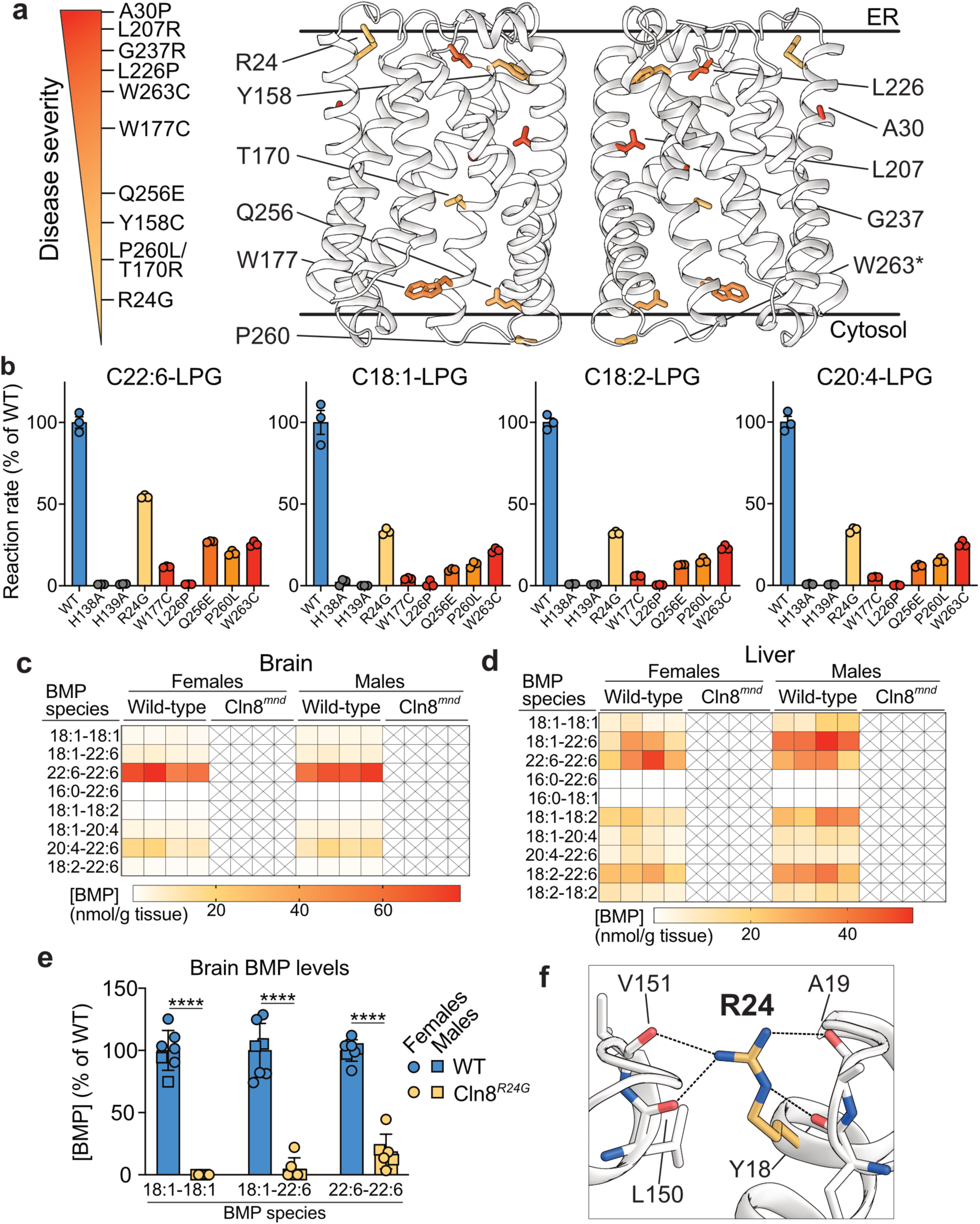
Impact of CLN8 mutations on enzymatic activity. a, Cartoon representation of the CLN8 homodimer with selected Batten disease missense mutations highlighted. Mutations are coloured by relative disease severity, classified based on reported age of onset and clinical symptoms. P260L and T170R were reported as compound heterozygous variants. W263 lies within the disordered C-terminal region not resolved in the structure. **b,** Residual activity of purified mutant CLN8 variants with the indicated acyl-CoA substrates, expressed relative to purified CLN8 WT, and coloured according to relative disease severity (n=3). **c- d,** Levels of the indicated BMP species in brains (**c**) and livers (**d**) of WT or Cln8*^mnd^* (*Cln8* frameshift mutation) mice (n = 4 WT and 4 homozygous males and females). No BMP species were detected in Cln8*^mnd^* tissues. **e,** Levels of the indicated BMP species in brains from 8- week-old WT and Cln8*^R24G^* homozygous knock-in mice, expressed as a percentage of the WT mean (n = 2 WT males and 2 homozygous males; 5 WT females and 5 homozygous females). **f,** Cartoon model showing electrostatic interactions between residue R24 and backbone carbonyl groups of neighbouring residues. In vitro assays are representative of at least two independent protein preparations. In f, ****P < 0.0001 by unpaired two-tailed Student’s t- test.

We purified six Batten disease-associated CLN8 mutants capable of restoring BMP levels, alongside wild-type (WT) CLN8 and catalytically inactive H138A and H139A active-site mutants (Extended Data Fig. 7a,b). In initial rate assays using mixed acyl-CoA, *S,S*-GPG, and purified proteins, all six disease mutants displayed markedly reduced activity relative to WT CLN8, whereas H138A and H139A were completely inactive (Fig. 3b). Similar results were obtained using ^14^C18:1-CoA as an acyl donor (Extended Data Fig. 7c).

The impact of each mutation on enzymatic activity broadly correlated with reported disease severity, with milder mutations tending to retain more activity than severe ones (Fig. 3a,b). However, exceptions emerged - namely, W263C causes late-infantile onset disease^32^, phenotypically analogous to a complete CLN8 loss-of-function, yet retained ∼25% of WT activity (Fig. 3b). This suggests that additional factors not investigated here, such as CLN8 protein stability, post-translational regulation in cells, or patient genetic background, may influence disease onset and progression.

To define the catalytic steps disrupted by each mutation, we monitored formation of the fluorescent NBD-C18:1-CLN8 intermediate. The L226P and W263C variants showed reduced fluorescence relative to wild type upon incubation with NBD-C18:1-CoA, consistent with a defect in the initial acylation (“ping”) step (Extended Data Fig. 7d). In contrast, the W177C and P260L mutants formed the intermediate similarly to wild-type CLN8 but showed only partial consumption in the presence of *S,S*-GPG, indicating impaired *S,S*-GPG binding or positioning in the active site (Extended Data Fig. 7d,e). The R24G and Q256E variants formed the intermediate with wild-type efficiency, and its levels similarly decreased in the presence of *S,S*-GPG, suggesting a slower rate of *S,S*-LPG product release in these mutants (Extended Data Fig. 7d,e).

To assess whether the enzymatic defects caused by CLN8 mutations translate into BMP deficiency in vivo, we analysed two complementary mouse models: Cln8*^mnd^*, a spontaneous *Cln8* frameshift allele widely used as a CLN8 Batten disease model^33–35^, and Cln8*^R24G^*, a knock-in allele leading to the R24G missense substitution that was engineered to model the later-onset CLN8 disease variant called Northern epilepsy, which is enriched in Finnish population^29,36^. Consistent with complete loss of CLN8 function, homozygous Cln8*^mnd^* mice displayed a complete loss of detectable BMP across brain, liver, and circulation (Fig. 3c,d; Extended Data Fig. 6d). Moreover, C22:6-LPG levels were profoundly reduced and GPG levels markedly elevated, supporting the expected substrate-product relationship (Extended Data Fig. 6e,f).

In homozygous Cln8*^R24G^* brains, BMP abundance was also markedly reduced in both sexes compared to WT controls (Fig. 3e). Notably, although the dominant C22:6-containing BMP species were still strongly reduced, they were relatively better preserved than other BMP species, consistent with the preferential retention of C22:6-CoA activity (∼50% of WT) compared to other acyl-CoA substrates (∼25%) for purified R24G mutant in vitro (Fig. 3b,e). Located at the TM1–TM4 interface and distant from the active site, R24 likely stabilises the CLN8 fold by mediating electrostatic interactions that tether TM1 to TM4 (Fig. 3f). We propose that, in R24G homozygous patients, these residual C22:6-containing brain BMP species sustain enough lysosomal function to delay disease onset, and that strategies aimed at stabilising BMP levels, such as inhibiting its degradation via PLA2G15^18^, may offer therapeutic benefit in patients where some CLN8 activity is retained.

Taken together, our biochemical and in vivo data show that CLN8 mutations cause a graded loss of acyltransferase activity that tracks with both tissue BMP levels and disease severity. Thus, impaired BMP synthesis emerges as the crucial biochemical defect underlying CLN8 Batten disease.

### *S,S*-LPG restores BMP synthesis in CLN8 deficiency

Finally, we tested whether supplementing *S,S*-LPG could restore BMP levels in CLN8-deficient models. Because *S,S*-LPG serves as both the acyl donor and acceptor for CLN5 during *S,S*- BMP synthesis, its supplementation is expected to yield a single symmetric BMP species in which both acyl chains match the supplemented *S,S*-LPG. We synthesised *S,S*-LPG containing a C18:1 acyl chain due to its relative chemical stability and because the resulting 18:1-18:1 BMP is abundant in both cultured cells and animal tissues (Fig. 4a). As a control, we used chemically synthesised *R,rac*-C18:1-LPG (an equal mixture of *R,R*-C18:1-LPG and *R,S*-C18:1- LPG) (Fig. 4a). This material is commercially available and chemically analogous to *S,S*-C18:1- LPG but carries the wrong stereochemistry to enter the CLN5-dependent step of the BMP synthesis pathway when CLN8 is absent.

**Fig. 4:**
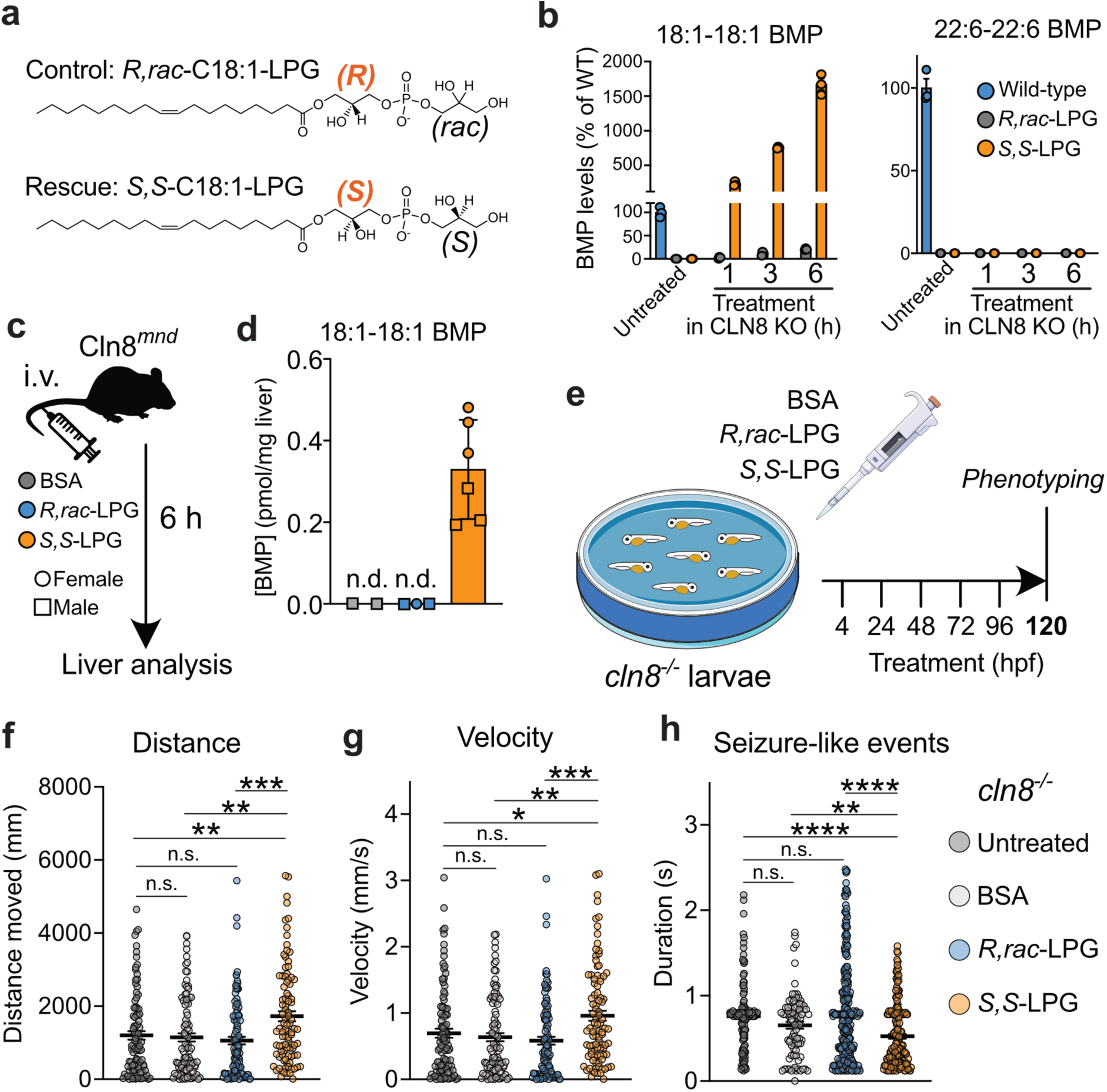
***S,S*-LPG restores BMP synthesis in CLN8-deficient models. a,** Chemically synthesised LPG stereoisomers used in rescue experiments. The rescue and control compounds differ only in the stereochemistry of the glycerol backbone. **b,** Levels of 18:1–18:1 and 22:6–22:6 BMP species in wild-type or CLN8 KO HeLa cells treated with 10 µM *R,rac*-LPG or *S,S*-LPG for the indicated times. Wild-type cells were untreated. **c,** Schematic of the intravenous LPG injection protocol in homozygous Cln8*^mnd^* mice (single bolus of 100 nmol BSA-conjugated lipid). **d,** Hepatic 18:1-18:1 BMP levels in Cln8*^mnd^* mice 6 h after injection with *S,S*-LPG (n = 3 males, 3 females), *R,rac*-LPG (n = 2 males, 1 female) or BSA vehicle (n = 2 males). BMP remained undetectable in BSA- and *R,rac*-LPG–injected mice**. e,** Schematic of the *cln8*^-/-^ zebrafish larvae treatment protocol (compound exposure at a final lipid concentration of 5 µM from 4 to 120 hpf with daily refresh; locomotor assays and local field potential recordings at 120 hpf). **f,g,** Locomotor activity (distance travelled and swimming velocity) recorded over 30 min in cln8^-/-^ larvae at 120 hpf (untreated n=97, BSA n=97, R,rac-LPG n=103, S,S-LPG n=95; 3 independent experiments). **h,** Duration of seizure-like events detected by local field potential recordings at 120 hpf (n=8 larvae per group). Statistical analysis in **(f–h)** was performed using the Kruskal–Wallis test followed by Dunn’s post-hoc test (*p < 0.05; **p < 0.01; ***p < 0.001; ****p < 0.0001).

In CLN8 KO HeLa cells, supplementation with BSA-conjugated *S,S*-C18:1-LPG, but not *R,rac*- C18:1-LPG, triggered a rapid and selective rise in 18:1-18:1 BMP, driving its levels well beyond those of untreated wild-type cells (Fig. 4b). As expected for the direct condensation of two *S,S*-LPG molecules by CLN5, other BMP species, including 22:6-22:6 BMP, remained undetectable, since CLN8 KO cells lack the capacity to introduce new acyl groups into *S,S*- GPG (Fig. 4b). Comparable increases in total cellular C18:1-LPG levels after both treatments confirmed that *S,S-* and *R,rac*-LPG were taken up with similar efficiency (Extended Data Fig. 8a). Notably, 18:1-18:1 BMP formation following *S,S*-C18:1-LPG supplementation was CLN5- dependent, as it was abolished in CLN5/8 DKO HeLa cells (Extended Data Fig. 8b). In these cells, C18:1-LPG accumulated to levels exceeding those in CLN8 KO cells, consistent with a block in its conversion to *S,S*-BMP (Extended Data Fig. 8c).

Next, we asked whether exogenous *S,S*-LPG could restore BMP synthesis in CLN8-deficient animals. To test this, we intravenously injected homozygous Cln8*^mnd^* mice with a bolus of BSA-conjugated *S,S*-LPG or *R,rac*-LPG (100 nmol/mouse) and quantified hepatic 18:1-18:1 BMP 6 h later (Fig. 4c). *S,S*-LPG administration led to the appearance of 18:1-18:1 BMP in the liver, whereas this species remained undetectable in vehicle- and *R,rac*-LPG–treated controls (Fig. 4d). Thus, exogenous *S,S*-LPG – the CLN8 reaction product – can bypass CLN8 deficiency and restore BMP synthesis in vivo.

Last, we investigated whether *S,S*-LPG could improve neurological phenotypes in CLN8- deficient animals. For this purpose, we chose *cln8* mutant zebrafish, whose larvae recapitulate key pathological signatures of CLN8 Batten disease – namely impaired motor function, seizure-like brain activity, and lysosomal dysfunction^37^. *Cln8*^-/-^ larvae were treated with LPG stereoisomers or vehicle by adding the compounds directly to the fish water from 4 to 120 hours post-fertilisation (hpf), and locomotor behaviour and seizure-like activity were assessed at 120 hpf (Fig. 4e). *S,S*-LPG treatment increased both distance travelled and swimming velocity compared with untreated, vehicle-treated, and *R,rac*-LPG-treated mutants (Fig. 4f,g). Local field potential recordings at the same stage similarly revealed shorter seizure-like events in *S,S*-LPG–treated larvae, whereas event power showed high variability across all treatment groups (Fig. 4h, Extended Data Fig. 8d). Together, these data show that *S,S*-LPG treatment in *cln8*^-/-^ zebrafish ameliorates key behavioural and electrophysiological deficits in vivo, supporting BMP deficiency as a central driver of pathology in CLN8 Batten disease and highlighting BMP precursor supplementation as a promising therapeutic strategy.

## Discussion

In this work, we identify CLN8 as a stereospecific GPG acyltransferase that catalyses the formation of *S,S*-2-acyl-LPG, a key intermediate in the *S,S*-2,2’-BMP synthesis pathway (Fig. 5). A separate study supports this conclusion by also independently identifying GPG as an intermediate in BMP biosynthesis^38^. Although CLN8 localises to the ER/Golgi, the final acyl transfer completing BMP synthesis occurs in the lysosomal lumen, thus requiring *S,S*-2-acyl- LPG trafficking between compartments (Fig. 5). This organisation means that CLN8 determines the acyl composition of mature BMP species during synthesis. To remodel BMP acyl tails, BMP must first be deacylated in the lysosome to *S,S*-GPG, exported back to the ER in a CLN3-dependent manner^39^, and then reacylated by CLN8 to re-enter the pathway (Fig. 5). Accordingly, CLN8 substrate selectivity and cellular acyl-CoA composition together shape BMP molecular diversity, with the enzyme’s strong preference for C22:6-CoA explaining the enrichment of C22:6-containing BMP species in the brain.

**Fig. 5:**
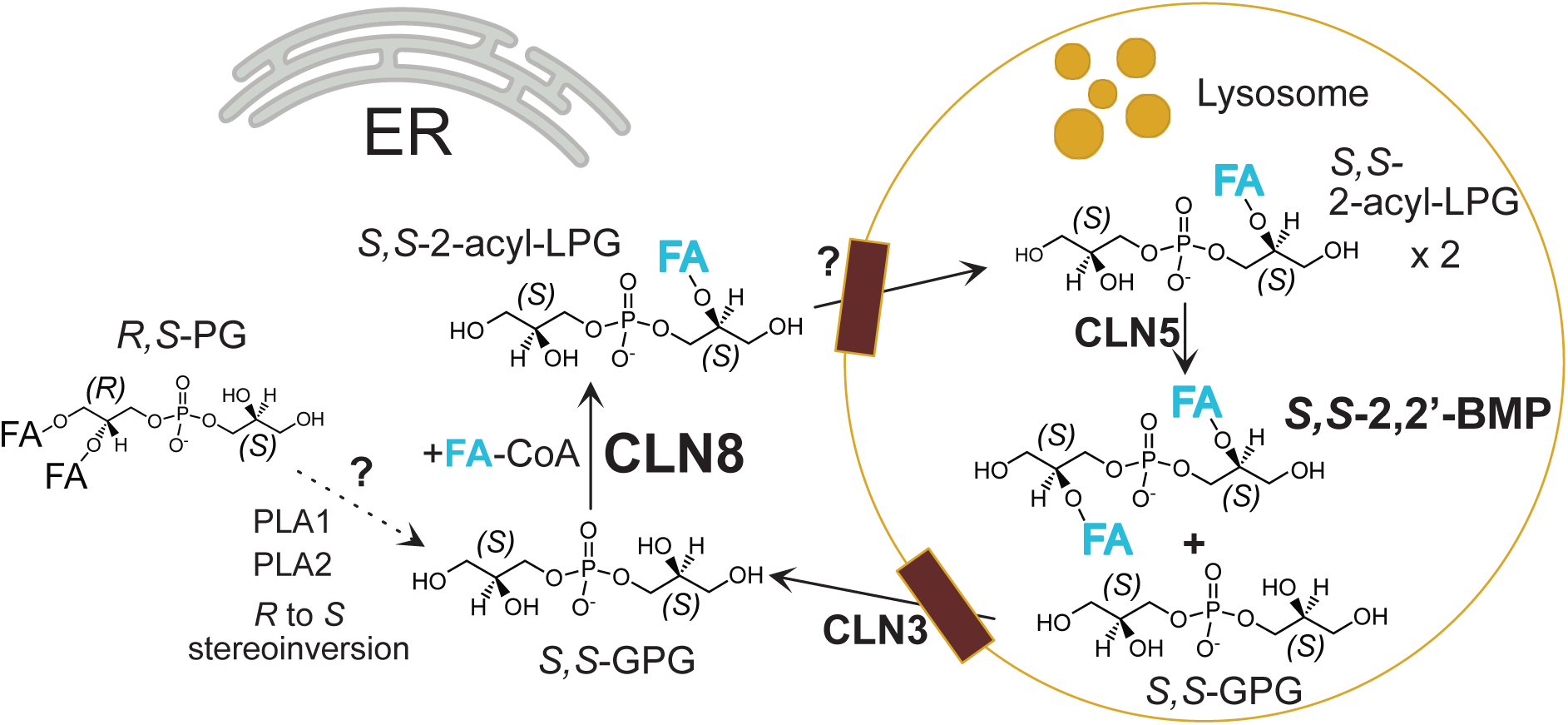
Working model of BMP biosynthesis. *R,S*-phosphatidylglycerol (PG) is first deacylated and undergoes *R*-to-*S* stereoinversion through an undefined mechanism to generate *S,S*- glycerophosphoglycerol (GPG). CLN8 then catalyses the key acyl transfer step, converting *S,S*- GPG to *S,S*-2-acyl-lysophosphatidylglycerol (LPG) at the endoplasmic reticulum (ER). *S,S*-2- acyl-LPG enters the lysosomal lumen via an unknown route, where CLN5 uses it as both acyl donor and acyl acceptor to complete *S,S-*2,2’*-*BMP synthesis, producing *S,S*-GPG as a by- product. This *S,S*-GPG is exported back to the ER in a CLN3-dependent manner, where it can be re-acylated by CLN8. BMP degradation by lysosomal lipases also yields *S,S*-GPG, which is similarly exported via CLN3 to re-enter the pathway. Because CLN8 generates the pool of *S,S*- 2-acyl-LPG used by CLN5, its enzymatic activity ultimately defines the acyl composition of BMP species. PLA – phospholipase A.

The structural and mechanistic insights presented here open opportunities for targeted modulation of BMP synthesis. The CLN8 paralog CERS6 is inhibited by the fungal toxin fumonisin B1, which acts as an acyl acceptor and traps itself within the enzyme cavity once acylated^19^. By analogy, our structural data enable the rational design of similar acyl- accepting inhibitors targeting the cytoplasm-facing catalytic cavity of CLN8. Such compounds would serve as valuable chemical tools to dissect the physiological roles of BMP in diverse contexts and could have therapeutic relevance in conditions where transient suppression of lysosomal catabolism is advantageous.

The essentiality of CLN8 for BMP synthesis is underscored by the complete loss of all BMP species in the commonly used Cln8*^mnd^* Batten disease mouse model. This striking phenotype is mirrored in CLN8 KO human cell lines^2^, highlighting the conserved and non-redundant role of CLN8 in BMP biosynthesis. Importantly, BMP synthesis can be restored by supplementing *S,S*-LPG – the CLN8 reaction product – in CLN8-deficient human cells and homozygous Cln8*^mnd^* mouse liver. Furthermore, this metabolic rescue improves the neurological phenotypes of CLN8-deficient zebrafish larvae, demonstrating that BMP deficiency is the primary driver of neuropathology in CLN8 Batten disease. Supporting this, Batten disease- causing CLN8 missense mutations reduce CLN8 enzymatic activity and BMP levels in proportion to clinical severity.

The therapeutic potential of *S,S*-LPG now justifies further rescue studies in mammalian CLN8-deficient models to optimise formulation, dosing, and delivery. More broadly, because additional Batten disease genes, including CLN3^40^ and CLN5, participate in BMP homeostasis, BMP deficiency may represent a unifying pathogenic mechanism across multiple Batten disease subtypes – a hypothesis that now demands systematic investigation.

## Methods

### Reagents and antibodies

Comprehensive lists of the antibodies and the chemicals/reagents used in this study are provided in Supplementary Tables 2 and 3, respectively.

### Plasmids

Details of plasmids used in this study are provided in Supplementary Table 4.

### Mouse studies

Tissues from age-matched (6-7 months old) wild-type and homozygous Cln8*^mnd^* mice were obtained from Sanford Research (Sioux Falls, SD, USA) for total BMP analysis. Animals were bred, housed, and euthanized under standardized conditions in an AAALAC International– accredited facility in accordance with approved Institutional Animal Care and Use Committee (IACUC) protocols (USDA license 46-R-0009; protocol #2023-0102). All procedures complied with institutional, federal, and international ethical guidelines governing the humane care and use of research animals.

Tissues from age-matched (8 weeks old) wild-type and homozygous Cln8*^R24G^* mice^36^ and their littermate controls were obtained from University of Oulu (Finland). Animal care was carried out with the support of the Oulu Laboratory Animal Centre Research Infrastructure, University of Oulu (Finland). All experimental procedures were conducted in accordance with the Finnish national legislation, EU Directive 2010/63/EU, and the ethical standards of University of Oulu. All animal procedures were approved by the Project Authorisation Board at the State Provincial Office of Southern Finland (ESAVI/33743/2019 and ESAVI/16903/2024).

The LPG rescue experiment complied with all relevant ethical regulations and was conducted in accordance with ARRIVE 2.0 guidelines. All procedures were reviewed and approved by the University of Cambridge Animal Welfare and Ethical Review Body (AWERB). Animal work and breeding was performed under the UK Animals (Scientific Procedures) Act 1986 (Home Office Project Licence PPL no. PP1740969) and in line with EU Directive 2010/63/EU. The Cln8*^mnd^* mouse line was obtained from The Jackson Laboratory, and homozygous experimental animals were generated by crossing heterozygous parents. Both female and male mice were used in accordance with the 3Rs principles. Mice were housed in individually ventilated cages (Tecniplast) at 20–24 °C and 45–65% humidity on a 12 h light– dark cycle, with ad libitum access to SAFE 105 universal diet (Safe Diets) and water.

For the LPG rescue experiment, custom-synthesised *S,S*-C18:1-LPG was dissolved in chloroform, dried under a stream of nitrogen to remove solvent, and the resulting lipid film reconstituted in 12% fatty acid- and endotoxin-free BSA (A8806, Sigma) prepared in 150 mM NaCl to a final concentration of 1 mM, using gentle heating and sonication. The control lipid (*R,rac*-C18:1-LPG; 858125, Avanti) was prepared in parallel using the same procedure.

Conscious mice received an intravenous tail vein injection of 100 µl of lipid–BSA complex (or the matched BSA vehicle in 150 mM NaCl) and were euthanised 6 h later by cervical dislocation. Livers were immediately snap frozen for lipid analysis.

### Zebrafish studies

Zebrafish experiments were performed using a previously characterised *cln8*^-/-^ zebrafish line^37^. Homozygous mutants were generated by intercrossing adult *cln8*^-/-^ males and females. Adult fish were maintained at 28 °C on a 14 h light / 10 h dark cycle at a density of ≤5 fish per litre. Embryos were collected and raised at 28.5 °C in egg water from ‘Instant Ocean’ sea salts (60 µg/mL) or E3 medium (292.2 mg NaCl, 12.6 mg KCl, 48.6 mg CaCl₂·2H₂O, 39.8 mg MgSO₄ per litre), prepared according to standard protocols^41^. Developmental stages were recorded as hours post-fertilisation (hpf). All zebrafish procedures were carried out in accordance with EU Directive 2010/63/EU and were approved by the Italian Ministry of Health (authorization n. 1090/2024-PR) under the oversight of the University of Pisa Animal Care and Use Committee.

For LPG rescue experiments, *S,S*-C18:1-LPG and *R,rac*-C18:1-LPG control were prepared as described above, but reconstituted in BSA at a final concentration of 10 mM. Both treatments, or equivalent amount of BSA control, were added directly to egg water at a final concentration of 5 µM, and the treatments were repeated every 24 hours until phenotypic analysis.

Larval motor behaviour was assessed as described^37^. Locomotor activity (distance travelled and velocity) was measured in *cln8*^-/-^ siblings at 120 hpf using the DanioVision system with EthoVision XT automated tracking.

Electrophysiological recordings were performed in the forebrain of 120 hpf larvae to assess seizure-like activity (epilepsy-like phenotype), following established procedures^42^.

### Molecular biology

Site-directed mutagenesis was performed using the Q5 Site-Directed Mutagenesis Kit (E0554S, New England Biolabs) on pcDNA3.1(+)-C-HA and pcDNA5/FRT/TO plasmids containing wild-type CLN8 inserts, following the manufacturer’s instructions. Resulting mutant constructs were verified by Sanger sequencing using CMVFpCDNA3 and BGHRev primers (for pcDNA3.1(+)-C-HA) or pCMVF and BGHRev primers (for pcDNA5/FRT/TO), performed by Source Genomics. In cases where pcDNA3.1(+)-C-HA constructs did not yield detectable CLN8 mutant protein expression, plasmid integrity and sequence were further validated by whole-plasmid Oxford Nanopore sequencing.

### Human cell culture

HeLa (ECACC 93021013) cell line was obtained from Culture Collections, UK Health Security Agency. The Flp-In T-REx 293 cell line was purchased from Thermo Fisher Scientific (R78007). All cell lines were maintained in DMEM (11965084, Thermo Fisher Scientific) supplemented with 10% fetal bovine serum (FBS; Sigma-Aldrich) at 37°C in 5% CO₂. Cells were passaged at ∼90% confluence using TrypLE Express (12605010, Thermo Fisher Scientific) and routinely tested for mycoplasma contamination.

For transient transfection, cells cultured in standard growth medium were transfected with FuGENE HD (E2311, Promega) following the manufacturer’s protocol. Briefly, for one well of a six-well plate, 20 μl Opti-MEM (31985062, Thermo Fisher Scientific), 0.4 μg plasmid DNA, and 1.2 μl FuGENE HD were mixed. Medium was replaced the next day, and cells were analyzed 48–72 hours post-transfection.

For lipid analyses, confluent cells in six-well plates were washed twice with 3 ml ice-cold PBS on ice, then scraped into 500 μl ice-cold PBS using an inverted 200 μl pipette tip. Cell suspensions were transferred to pre-chilled 2-ml plastic tubes (3469-11, Thermo Fisher Scientific) and centrifuged at 500g, 4°C for 5 min. Supernatants were removed, and cell pellets were snap-frozen and stored at −70°C. Each well provided one replicate for lipid analysis.

For d5-PG labelling, 16:0-18:1 d5-PG (860385, Avanti) was dissolved in ethanol to prepare a 10 mM stock solution. The stock was added dropwise to complete DMEM to achieve a final concentration of 25 μM and mixed thoroughly by vortexing. Labelling was initiated by replacing the culture medium with the prewarmed d5-PG-containing medium.

For LPG rescue experiments, *S,S*-C18:1-LPG and *R,rac*-C18:1-LPG control were prepared as described above, but reconstituted in BSA at a final concentration of 10 mM. Both treatments were added directly to cell culture medium at a final concentration of 10 µM, and cells were harvested after the incubation times indicated in the figure legends.

### Human knockout cell line generation

HeLa knockout (KO) cell lines were generated by transfecting cells with recombinant complexes of Cas9 protein and synthetic guide RNAs (gRNAs). The HeLa CLN8 KO and corresponding control cell lines were previously established as described^2^. To generate a matched set of control, CLN8 KO, CLN5 KO, and CLN5/8 double KO (DKO) lines, both the published control and CLN8 KO cells were edited again using either a non-targeting or a CLN5-targeting CRISPR strategy, resulting in four parallel cell lines used in this study.

For transfection, cells were seeded in six-well plates in standard growth medium and transfected the following day at approximately 30% confluence. Transfections were performed using Lipofectamine CRISPRMAX Cas9 Transfection Reagent (CMAX00001, Thermo Fisher Scientific), TrueCut Cas9 Protein v2 (A36497, Thermo Fisher Scientific), and custom-synthesized TrueGuide synthetic gRNAs (Thermo Fisher Scientific), following the manufacturer’s instructions. Each CLN5 knockout transfection contained 6250 ng Cas9 protein and 600 ng of each targeting gRNA (GAAAUGUACAGUUGCCAAGU and GCCUUGAUUACACCAGAAAG), while control transfections contained 6250 ng Cas9 protein and 1200 ng of non-targeting gRNA (A35526, Thermo Fisher Scientific) to generate the corresponding control cell lines.

After transfection, the medium was replaced after 24 h, and cells were passaged into 10-cm dishes the following day. Control and KO populations were subsequently expanded in multiple 15-cm dishes, cryopreserved, and later thawed for validation by Western blotting and quantitative proteomics. All experiments were performed using low-passage cells following thawing.

### Human Flp-In T-REx 293 inducible cell line generation

Flp-In T-REx 293 cells were seeded in six-well plates and transfected at ∼25% confluence. Transfections were performed using a mixture of 20 μl Opti-MEM, 0.36 μg pOG44 plasmid, 0.04 μg pcDNA5/FRT/TO plasmid encoding CLN8 WT or mutant, and 1.2 μl FuGENE HD. Medium was replaced after 24 hours, and cells were cultured for an additional 48 hours before passaging into 6-cm dishes. Stable integrants were selected in standard medium containing 50 μg/ml hygromycin B for 2 weeks, with passaging upon confluence.

Hygromycin-resistant pools were expanded, aliquoted, and cryopreserved. An aliquot was thawed and tested for inducible protein expression by tetracycline dose–response (48 h) and time-course assays (at the optimized dose). All CLN8 WT and mutant proteins were purified following induction with 1 μg/ml tetracycline for 48-72 hours.

### Yeast strain generation for protein purification

*Saccharomyces cerevisiae* strain W303.1B (MATα leu2-3,112 trp1-1 can1-100 ura3-1 ade2-1 his3-11,15) (ATCC: 201238) was used for protein expression and purification. Yeast line expressing human CLN8 was generated according to the published protocol^43^. Briefly, pYES2/CT C-HIS-CLN8 plasmid was transformed into competent W303.1B cells by lithium acetate/polyethylene glycol and heat-shock. Transformants were selected on SC-Ura + 2% glucose agar plates at 30°C for 48–72 h. Individual colonies were inoculated into 2 ml SC-Ura + 2% glucose liquid medium and grown overnight at 30°C with shaking at 225 rpm. Glycerol stocks were prepared by mixing 750 μl of culture with 750 μl of 30% glycerol and stored at −70°C.

### CLN8 protein purification from human cells

Flp-In T-REx 293 cells with inducible expression of C-terminal FLAG-STREP-tagged CLN8 WT or mutant variants were seeded in 15-cm dishes (10 dishes per typical purification) in standard culture medium. The following day, at ∼25% confluence, expression was induced with 1 μg/ml tetracycline for 48-72 hours. Cells were then washed once with ice-cold PBS, scraped into ice-cold PBS on ice, and pelleted by centrifugation at 500g, 4°C for 5 min.

Pellets were stored at −70°C until use.

For purification, frozen pellets were thawed and resuspended in Hepes buffer [20 mM Hepes (pH 7.4), 150 mM NaCl]. Solubilization was performed by adding 10% LMNG (Anatrace) to a final concentration of 1.5%, followed by incubation at 4°C for 2 hours. Lysates were cleared by centrifugation at 128,000g, 4°C for 1 hour. Meanwhile, 600 μl Strep-Tactin XT 4Flow bead slurry (Iba) was washed twice with Milli-Q water and once with wash buffer [20 mM Hepes (pH 7.4), 150 mM NaCl, 0.1% LMNG]. The clarified supernatant was incubated with the washed beads for 2 hours at 4°C. Beads were then washed thrice with wash buffer to remove unbound proteins. CLN8 was eluted in 1 ml elution buffer [50 mM Hepes (pH 7.4), 150 mM NaCl, 0.05% LMNG, 100 mM biotin]. Excess biotin was removed by passage through PD Minitrap G-25 columns (17085101, Cytiva). Protein concentration was determined using a NanoDrop ND-8000 at 280 nm. Purified proteins were aliquoted and stored at −70°C. All CLN8 mutants were purified using the same protocol alongside WT controls.

### CLN8 protein purification from yeast

W303 yeast strain expressing codon optimised C-His-tagged human CLN8 was retrieved from glycerol stock and scaled up to 10 litres in YPG + 0.1% glucose at 30°C with shaking (225 rpm). Expression was induced by adding galactose to a final concentration of 0.4% for 4 h.

Cells were harvested by centrifugation, washed with water, and stored at −20°C.

For membrane preparation, cell pellets were thawed and homogenized in breaking buffer [0.65 M sorbitol, 100 mM Tris-HCl (pH 8.0), 5 mM EDTA, 5 mM amino hexanoic acid, 5 mM benzamidine hydrochloride]. Cells were lysed using a Constant Systems Cell Disruptor. Debris was cleared by centrifugation (3,000g, 20 min, 4°C), and membranes were pelleted by ultracentrifugation (125,000g, 1 h, 4°C). The membrane pellet was washed in wash buffer [0.65 M sorbitol, 100 mM Tris-HCl (pH 7.4), 5 mM amino hexanoic acid, 5 mM benzamidine hydrochloride], pelleted again, then washed in TBG buffer [100 mM Tris-HCl (pH 7.4), 10% glycerol] and pelleted once more. The final membrane pellet was weighed, resuspended in TBG, aliquoted, and stored at −70°C.

For purification, yeast membranes were solubilized in 21 ml buffer containing 1.5% LMNG, 20 mM Hepes (pH 7.0), 150 mM NaCl, 30 mM imidazole, and protease inhibitors, and incubated for 1.5 h at 4°C. Insoluble material was removed by centrifugation (118,000g, 60 min), and the supernatant was collected. Ni-Sepharose excel resin (Cytiva) was equilibrated with Milli-Q water and wash buffer A [20 mM Hepes (pH 7.0), 150 mM NaCl, 40 mM imidazole, 0.1% LMNG]. The soluble fraction was incubated with resin for 2h at 4°C. Resin was washed thrice with wash buffer A and once with wash buffer B [20 mM Hepes (pH 7.0), 100 mM NaCl, 2.5mM CaCl2 and 0.05% LMNG]. Bound proteins were resuspended in 1.5 ml buffer B, and elution was performed by overnight incubation at 4°C with 20 μg Factor Xa protease and 2.5 mM CaCl₂. Protein concentration was determined by measuring absorbance at 280 nm (NanoDrop ND-8000).

### Size exclusion chromatography (SEC)

CLN8 was purified from yeast as described above. The protein was concentrated to 3.5 mg ml⁻¹ using a 10 kDa molecular-weight cut-off (MWCO) centrifugal concentrator (UCF501008, Millipore). A 130 µl aliquot was injected onto an ÄKTA Purifier system (GE Healthcare) equipped with a Superdex 200 Increase 10/300 GL column (GE Healthcare) equilibrated in SEC buffer [50 mM Hepes (pH 7.4), 150 mM NaCl, 0.05% LMNG]. Fractions corresponding to the main elution peak (indicated by dotted lines) were pooled and reconcentrated using the 10 kDa MWCO concentrator. The resulting sample was analysed by SDS–PAGE and in vitro activity assays, and the purified protein was used for cryo-EM grid preparation.

### Sample preparation and cryo-EM data acquisition

Cryo-EM grids were prepared using CLN8 at three concentrations (1, 2, and 3 mg ml⁻¹). For the apo condition, the protein was incubated with 0.05% (w/v) fluorinated octyl maltoside for 45 min on ice before grid preparation. For the substrate-bound conditions, CLN8 (2 mg ml⁻¹) was incubated in SEC buffer containing 100 µM C18:1-CoA or 100 µM C22:6-CoA for 1 h at room temperature, followed by addition of 0.05% fluorinated octyl maltoside and incubation for 45 min on ice. For each condition, 3 µl of sample were applied to glow- discharged Quantifoil R1.2/1.3 300-mesh holey carbon copper grids (30 s at 10 mA) and vitrified in a Vitrobot Mark IV (Thermo Fisher Scientific) operated at 4 °C and 95% relative humidity (blot time = 3.0 s; blot force = −10).

Datasets were collected from a single grid per condition on a Titan Krios (Thermo Fisher Scientific) operating at 300 kV and nominal magnification ×165,000, equipped with a Falcon 4i detector and a Selectris X energy filter in counting mode. Movies were recorded in EPU v3.1 at a physical pixel size of 0.729 Å pixel⁻¹, with a target defocus range of −1.8 to −2.4 µm in 0.2-µm increments; autofocus was executed every 10 µm of stage travel. The total exposure was 49.81, 49.89 and 50.14 e⁻ Å⁻² for the apo, C18:1-bound, and C22:6-bound datasets, respectively, with dose rates of 6.03, 6.04 and 6.07 e⁻ pixel⁻¹ s⁻¹. Data were acquired using aberration-free image shift (AFIS) with four exposures per hole, a 5-s delay after stage shifts, and a 1-s delay after image shifts.

### Cryo-EM data processing

Processing was performed in cryoSPARC v3.3.2. For each dataset, ∼9,000–10,000 movies were collected. Movie-frame alignment, contrast transfer function (CTF) estimation, particle picking, and particle extraction (box size, 416 pixels) were carried out in Warp^44^. Particles were initially Fourier-binned to 2.35 Å/pixel and subjected to multiple rounds of 2D classification to remove artefacts, contaminants, and low-quality particles. Ab initio reconstruction in cryoSPARC produced six initial classes, which were used for heterogeneous refinement to separate particle populations (e.g., CLN8 dimers vs empty micelles). Two successive heterogeneous refinements were performed, after which selected particles were re-extracted at the full pixel size (0.729 Å/pixel) and refined using non-uniform refinement with the best heterogeneous-refinement map as reference, yielding final reconstructions at 2.69–3.65 Å resolution. Cryo-EM data collection, refinement and validation statistics are summarised in Supplementary Table 1.

For the CLN8 apo dataset, an additional local refinement was performed after non-uniform refinement. A mask was generated in UCSF Chimera v1.13.1, and the final map was Gaussian-filtered using “vop gaussian” (σ = 4); the threshold was adjusted to encompass the dimeric assembly.

### Model building, validation, and structural analysis

AlphaFold-predicted CLN8 models^45,46^ were rigid-body fitted into the cryo-EM density in UCSF Chimera^47^, with side chains pruned to match the density. The model was rebuilt using ISOLDE^48^ within ChimeraX^49^ and further refined in Coot^50^. For the CLN8 + C18:1-CoA, CLN8 + C22:6-CoA, and CLN8-apo CoA-bound datasets, clear densities were present in the CLN8 structures; the corresponding ligands were modelled with ChemEM^51^ guided by the cryo-EM maps. Restraint files for C18:1, C22:6, and CoA were taken from the CCP4 monomer library. Models were validated with MolProbity^52^. Final positional minimisation and group B-factor refinement were performed in Servalcat^53^ against the cryoSPARC-refined maps. Structural figures were prepared in PyMOL (Schrödinger, LLC).

### Denaturing PAGE, in-gel fluorescence, Coomassie staining, and Western blotting

Twelve percent SDS–polyacrylamide gels were manually cast using Invitrogen SureCast Acrylamide Solution (40%). Samples were prepared at room temperature without heating, using Pierce Lane Marker Reducing Sample Buffer at a final concentration of 2.5x. Notably, heating led to high-molecular weight aggregation of CLN8 proteins, consistent with observations for other transmembrane proteins^54^. Electrophoresis was performed in Tris-glycine-SDS buffer at 120 V for stacking and 180 V for resolving gels.

For Coomassie staining, gels were incubated in 15 ml InstantBlue Protein Stain (Abcam) overnight, then destained with water overnight. Stained gels were imaged using a flatbed scanner.

For in-gel NBD fluorescence, 1.5 μg purified protein was incubated with 5 μM NBD-C18:1- CoA alone or with 5 μM NBD-C18:1-CoA plus 500 μM S,S-GPG at 37°C for 15 min. Reactions were stopped by adding sample buffer (2.5x), then analysed by SDS-PAGE as above.

Fluorescent images were acquired using an Amersham Typhoon imager (Cy2 setting), followed by Coomassie staining and imaging.

For Western blotting, proteins were transferred onto 0.45 μm PVDF membranes (Thermo Fisher Scientific) in chilled Tris-glycine buffer with 10% methanol at 90 V for 100 min using a Mini Trans-Blot system (Bio-Rad) with an ice pack. Membranes were blocked in 5% skimmed milk in TBST, then incubated overnight at 4°C with primary antibodies (Supplementary Table 2). After washing, membranes were incubated with secondary antibodies (1:5000) for 1 hour at room temperature. Detection was performed by enhanced chemiluminescence (Thermo Fisher Scientific) and imaged on a ChemiDoc MP system (Bio-Rad). Where indicated, blots are shown as overlays of chemiluminescence and colorimetric images to display molecular weight markers.

### Quantitative proteomics

Cell pellets were lysed in 100 µl lysis buffer consisting of 1% sodium deoxycholate (SDC; Sigma, #D6750), 100 mM triethylammonium bicarbonate (TEAB; Sigma, #T7408), 10% isopropanol, 50 mM NaCl, 1× protease and phosphatase inhibitor cocktail (Thermo Fisher Scientific, #78441) and nuclease (Thermo Fisher Scientific, #88700; 0.5 µl per 1 ml buffer). Lysates were briefly sonicated to improve sample dissolution, and protein concentrations were determined using the Quick Start Bradford Protein Assay (Bio-Rad) according to the manufacturer’s instructions. For each sample, 10 µg protein was reduced and alkylated in 5 mM tris(2-carboxyethyl)phosphine (TCEP; Thermo Fisher Scientific, #77720) and 10 mM iodoacetamide at room temperature for 1 h in the dark, followed by overnight digestion at room temperature with 500 ng trypsin (Pierce, #90058). Peptides were acidified with trifluoroacetic acid (TFA) to precipitate SDC, which was removed by centrifugation at 12,000 g. The clarified peptide solutions were desalted using C18 Spin Tips (Pierce). Briefly, C18 tips were conditioned with 80% acetonitrile (Fisher Scientific) / 0.1% TFA and equilibrated with 0.1% TFA, after which acidified peptides were loaded, washed with 0.1% TFA, and eluted with 80% acetonitrile / 0.1% TFA. Eluates were dried in a SpeedVac (Eppendorf) and resuspended in 40 µl 0.1% formic acid; 2 µl (approximately 500 ng peptide) were injected for LC–MS analysis.

Peptides were analysed on a Vanquish Neo UHPLC system (Thermo Fisher Scientific) coupled to an Orbitrap Fusion Lumos Tribrid mass spectrometer (Thermo Fisher Scientific). Peptides were first trapped on a 300 µm × 5 mm C18 trap cartridge (5 µm, 100 Å), then separated over a 100-min gradient on a 75 µm × 50 cm C18 reversed-phase column (2 µm, 100 Å) at a flow rate of 300 nl min⁻¹. In each acquisition cycle, one full MS scan (m/z 375–1200) was acquired in the Orbitrap at 60,000 resolution with a normalised automatic gain control (AGC) target of 300% and automatic maximum injection time (MIT). Data-independent acquisition (DIA) spectra were then acquired in the Orbitrap at 30,000 resolution (normalised AGC 800%, automatic MIT, HCD collision energy 30%) using 42 DIA windows across 400-900 m/z precursor scan range.

Raw data were processed using DIA-NN (version 2.0). An in silico–predicted spectral library was generated from a human FASTA database (9606_2025_01_23) and used to analyse the DIA data, allowing one missed cleavage, N-terminal methionine excision, and carbamidomethylation of cysteine as a static modification. The peptide length range was set to 7–30 amino acids, precursor charge range to 1–4, precursor m/z range to 375–1200, and fragment ion m/z range to 145–1450. Match-between-runs was enabled, and the quantification strategy was set to “QuantUMS (high precision)”. MS1 and MS2 mass accuracy and retention time window (scans) were set to 5, 15 and 8, respectively. All other DIA-NN settings were left at default, using retention time–dependent cross-run normalisation.

### Chemical synthesis of *R,S*-GPG standard

The target compound *R,S*-GPG ((*R*)-2,3-dihydroxypropyl ((*S*)-2,3-dihydroxypropyl) hydrogen phosphate) was synthesized through a nine-step sequence starting from (*S*)-(2,2-dimethyl- 1,3-dioxolan-4-yl) methanol. In the first step, benzylation with benzyl bromide in DMSO using sodium hydride afforded (*S*)-4-((benzyloxy)methyl)-2,2-dimethyl-1,3-dioxolane (92% yield). Subsequent acidic hydrolysis (70% acetic acid, 65°C) yielded (*R*)-3-(benzyloxy) propane-1,2-diol (81%). Selective tritylation of the primary alcohol with trityl chloride, triethylamine, and DMAP in THF at 80°C gave (*S*)-1-(benzyloxy)-3-(trityloxy) propan-2-ol (81%). Further benzylation with benzyl bromide under basic conditions produced (*S*)-((2,3- bis(benzyloxy) propoxy) methanetriyl) tribenzene (86%).

Cleavage of the trityl group by heating in 70% acetic acid provided (*R*)-2,3- bis(benzyloxy)propan-1-ol (82%), which was phosphorylated with diphenyl phosphonate in pyridine to afford triethylammonium (*S*)-2,3-bis(benzyloxy)propyl phosphonate (59%). This intermediate was coupled with (*S*)-(2,2-dimethyl-1,3-dioxolan-4-yl) methanol using pivaloyl chloride activation, followed by oxidative iodine treatment, to yield triethylammonium (*S*)- 2,3-bis(benzyloxy)propyl (((*R*)-2,2-dimethyl-1,3-dioxolan-4-yl)methyl) phosphate (22%).

Acidic deprotection (TFA) furnished (*S*)-2,3-bis(benzyloxy)propyl ((*R*)-2,3-dihydroxypropyl) hydrogen phosphate (53%). Finally, hydrogenolysis over Pd(OH)₂ in MeOH delivered the target diol phosphate *R,S-*GPG in 10% yield.

All intermediates were characterized by MS and ¹H-NMR spectroscopy. Purifications were performed by silica or neutral alumina chromatography with appropriate solvent gradients.

### Synthesis of GPG stereoisomers

*R,R*-, *R,S*-, and *S,S*-GPG stereoisomers were generated by mild methanolysis from *R,R*-BMP (857136, Avanti), *R,S-*BMP (857133, Avanti), and *S,S*-BMP (857135, Avanti), respectively. 5 mg of each BMP were dissolved in 1 ml of 0.1 M NaOH in methanol and incubated at 37°C for 1 hour. Reactions were neutralized with 1 M acetic acid. Fatty acid methyl esters, residual BMP, and partially hydrolysed LPG contaminants were removed by butanol–methanol extraction, and the aqueous phase containing GPGs was collected and dried under nitrogen. The dried GPGs were resuspended in methanol, insoluble salts were removed by paper filtration, and the filtrate was again dried under nitrogen. Purified GPGs were dissolved in water, aliquoted, and stored at −20°C. Concentrations were determined by LC-MS/MS against a chemically synthesized *R,S*-GPG standard. LC–MS/MS analysis confirmed the absence of residual BMP or LPG in all GPG preparations.

### Chemical synthesis of *S,S*-C18:1-LPG

*S,S*-configured lysophosphatidylglycerol (18:1-(*S,S*ʹ)-LPG) was synthesized via a multi-step stereospecific route starting from (*R*)-(2,2-dimethyl-1,3-dioxolan-4-yl)methanol. The alcohol was first coupled with oleoyl chloride in dichloromethane containing pyridine and DMAP to yield (*R*)-(2,2-dimethyl-1,3-dioxolan-4-yl)methyl oleate. Subsequent acetal hydrolysis with aqueous acetic acid afforded (*R*)-2,3-dihydroxypropyl oleate, which was protected as the bis- tert-butyldimethylsilyl (TBS) ether using TBSCl and imidazole in THF/DMF. Selective desilylation with pyridinium hydrofluoride produced the mono-TBS derivative, (*R*)-2-((tert-butyldimethylsilyl)oxy)-3-hydroxypropyl oleate.

The free hydroxyl group was phosphitylated using 2-cyanoethyl (((*S*)-2,2-dimethyl-1,3- dioxolan-4-yl)methyl) diisopropylphosphoramidite and 1H-tetrazole in THF to generate a phosphite intermediate, which was oxidized with tert-butyl hydroperoxide to the corresponding phosphate triester. Removal of the cyanoethyl protecting group with triethylamine yielded the monoester phosphate, and final global deprotection with aqueous acetic acid (50 °C, 2 h) removed both the TBS and acetonide groups to yield 18:1-(*S,S*ʹ)-LPG. The product was neutralized with ammonium hydroxide and purified by preparative reverse- phase HPLC (Waters XBridge BEH C18, 10 mM NH₄HCO₃/ACN, 20–60% gradient over 15 min).

The final compound was obtained as an off-white gum (25% yield, 97% purity) and confirmed by ^1^H and ^31^P NMR (CD₃OD) and LC–MS/MS, consistent with the expected structure of 18:1-(*S,S*ʹ)-LPG.

### Synthesis of C18:1-LPG acyl isomer standards

*R,rac-*LPG (858125, Avanti) was used as the 1-C18:1-LPG standard. The 2-C18:1-LPG standard was generated as previously described^55^. Briefly, 1 mg of 18:1–18:1 PG (840475, Avanti) was incubated at 37 °C for 80 min in a reaction mixture containing 78 mM phosphate, 10% diethyl ether, 2 mM CaCl₂, 0.05% Triton X-100, and 4.5 µg mL⁻¹ *Rhizomucor miehei* lipase (A225779, Antibodies.com), which exhibits intrinsic PLA₁ activity (pH 7.4). The reaction was quenched by adding nine volumes of acidic methanol (pH 4.0). Both *R,rac*-LPG and enzymatically generated 2-C18:1-LPG were diluted to 1 µM in acidic methanol and analysed in parallel by LC–MS/MS.

### Acyltransferase activity assays

Enzymatic reactions were performed in 25 μl volumes in 1.5-ml microcentrifuge tubes using Hepes buffer [20 mM Hepes (pH 7.4), 150 mM NaCl, 0.05% LMNG]. Unless otherwise indicated, reactions contained 3.4 nM CLN8, 100 μM acyl-CoA (substrate A), and 500 μM LPG or GPG (substrate B). When using acyl-CoA mixtures, each was present at 100 μM. Reactions were assembled on ice and incubated at 37°C for 15 min (or as specified for time- course experiments). Reactions were quenched by adding either 500 μl butanol/methanol (3:1) for TLC or 200 μl 80% acetonitrile with 0.2% formic acid for LC-MS/MS.

For Km determination of *S,S*-GPG, acyl-CoA was fixed at 100 μM. Radiolabelling assays used a 10× tracer mix prepared by combining oleoyl-[1-¹⁴C]-CoA (American Radiochemicals) with unlabelled oleoyl-CoA at a 1:12.5 ratio, yielding a final acyl-CoA concentration of 100 μM in the reaction.

All assays included controls lacking enzyme, substrate A, or substrate B, run under identical conditions. For time-course assays, controls were incubated for the longest time point; for concentration series, controls contained the highest substrate concentration.

### Lipid extraction

Total lipids were extracted from cells, in vitro assays, and animal tissues using the butanol-methanol (BUME) method^56^. Extractions were performed in 2-ml screw-cap plastic tubes (3469-11, Thermo Fisher Scientific), with blank extractions run in parallel to monitor plastic- derived contaminants. For tissues, pre-weighed frozen samples were homogenized with 1.4- mm ceramic beads (19-645-3, Cole Parmer UK) directly in 0.5 ml ice-cold butanol:methanol (3:1). Subsequently, 0.5 ml of 1% acetic acid and 0.5 ml heptane:ethyl acetate (3:1) were added, and samples were vortexed vigorously for 5 min. After centrifugation (6,000g, 5 min), the upper organic phase was transferred to glass vials. A second extraction was performed on the remaining aqueous phase, and organic layers were combined. Solvents were evaporated under nitrogen, and dried lipid extracts were stored at −70°C until analysis.

### Glycerophosphoglycerol (GPG) extraction and LC-MS/MS quantification

Glycerophosphoglycerol (GPG) was extracted from mouse tissue by homogenising samples in 2-ml screw-cap tubes (3469-11, Thermo Fisher Scientific) containing 1 ml of 80% methanol and 1.4-mm ceramic beads (19-645-3, Cole Parmer UK). Homogenates were clarified by centrifugation (20,000 g, 15 min, 4 °C), and the resulting supernatants were transferred to glass vials for analysis.

GPG was analysed by LC–MS/MS using a Shimadzu 8060 triple quadrupole mass spectrometer (Shimadzu, UK). For initial MS¹ and product-ion scans, 2 µL of a 100 nM *R,S*- GPG standard in methanol was introduced by flow-injection (FIA), and spectra were acquired over an m/z range of 100–1000. Chromatographic separation was performed on an Atlantis Premier BEH Z-HILIC column (1.7 µm, 2.1 × 150 mm; Waters) using mobile phase A (15 mM ammonium acetate with 0.1% ammonium) and mobile phase B (100% methanol). The column was operated at a flow rate of 0.2 mL min⁻¹, and 2 µL of sample was injected.

HPLC was conducted on an LC-30AD system at 0.1 mL min⁻¹, with samples injected at 0.5 min using a SIL-30AC autosampler. The total run time was 6 min, with the following gradient: 0–1 min, 80% B; 1–3 min, 80%→20% B; 3–5 min, 20% B; 5–6 min, 80% B. No internal standard was used.

Source parameters were as follows: nebulising gas, 3 L min⁻¹; heating gas, 10 L min⁻¹; interface temperature, 300 °C; desolvation line (DL) temperature, 245 °C; heat-block temperature, 400 °C; drying gas, 10 L min⁻¹. GPG was detected in negative-ion mode using the following transitions: m/z 244.5→152.9 (dwell 100 ms; Q1 20 V; CE 15 eV; Q3 20 V), 244.5→78.95 (dwell 100 ms; Q1 20 V; CE 35 eV; Q3 20 V), and 244.5→170.95 (dwell 100 ms; Q1 20 V; CE 17 eV; Q3 21 V).

### Thin layer chromatography

Lipids extracted from in vitro assays were dissolved in 25 μl HPLC-grade chloroform and spotted at the base of 10 × 10 cm silica TLC plates (821050, Macherey-Nagel). Plates were developed in sealed glass chambers using a solvent system of chloroform:methanol:ammonium hydroxide (65:25:4, v/v) until the solvent front reached ∼1 cm from the top. Developed plates were dried under a laminar flow hood. For fluorescence detection, plates were imaged on an Amersham Typhoon imager using the Cy2 fluorescence setting. For phosphorimaging, TLC plates were exposed to pre-erased phosphor screens (GE Healthcare) in light-tight cassettes overnight, then scanned on the Typhoon imager with the phosphor imaging method. The migration of LPG and BMP reaction products was confirmed by *R,rac*-LPG (858125, Avanti) and *R,S*-BMP (857133, Avanti) standards, visualised by primuline staining and UV detection.

### Targeted LC–MS/MS analysis of lipids

Targeted quantification of LPGs and BMPs was performed on a Xevo TQ-S triple quadrupole mass spectrometer (Waters, UK) coupled to an I-Class ACQUITY UPLC system (Waters, UK). Samples were stored in glass vials at 8 °C prior to analysis, and 10 µL was injected via autosampler for each run. Chromatographic separation was achieved using a BEH C18 column (1 mm × 50 mm, 1.7 µm particle size, 130 Å pore size; Waters, UK) maintained at 30°C and fitted with a 0.2 µm UPLC in-line filter.

Mobile phase A consisted of 5% acetonitrile (ACN) and 0.1% formic acid (FA), and mobile phase B of 95% ACN and 0.1% FA, delivered at a flow rate of 0.2 mL min⁻¹ under the following gradient: 0–1.5 min, 5% B; 1.5–3 min, 5–40% B; 3-12 min, 40-70% B; 12-15 min, 70-100% B; and from 15 min to the end of the run, 100% B. The total run time was 20 minutes when measuring LPG alone, or 60 minutes when measuring both LPG and BMP.

Mass spectrometric detection was carried out in positive ion mode using multiple reaction monitoring (MRM) with electrospray ionization. The source parameters were as follows: capillary voltage, 3.1 kV; desolvation temperature, 500 °C; and source temperature, 150 °C. Nitrogen served as the curtain gas and argon as the collision gas. Cone voltages, collision energies, and analyte-specific MS/MS transitions are listed in Supplementary Table 5.

Quantification was performed in MassLynx 4.1 software using automatic peak selection followed by manual inspection and correction when required. LPGs and BMPs were quantified against a five-point calibration curve generated from 10-fold serial dilutions of *R,rac*-LPG (858125, Avanti) and *R,S*-BMP (857133, Avanti) standards, analysed alongside each sample batch. Multiple transitions were included for peak validation as both transitions should have identical retention times.

LPG species detected by LC–MS/MS were consistently resolved as two distinct peaks separated by 0.2–0.5 min in retention time, corresponding to the 2-acyl- and 1-acyl-LPG positional isomers, with the 2-acyl species eluting earlier under reverse-phase conditions^57^. In all CLN8 in vitro assays, only the first (2-acyl) LPG peak was detected and quantified; for the C18:1 species, the identity of this peak was confirmed using 1-acyl and 2-acyl C18:1-LPG standards. In biological samples, where appropriate, the integrated areas of both positional acyl isomers were combined to represent the total abundance of the corresponding LPG species.

### Statistical analysis, data visualisation, and reproducibility

All data are presented as means with error bars representing the SEM. The number of replicates is indicated in the corresponding figure legends. Where applicable, data are shown as fold changes relative to a specified control, as detailed in the legend. Each data point from in vitro assays represents a single measurement. Statistical methods are specified in the figure legends. Comparisons between two groups were performed using a Student’s t test; for more than two groups, a one-way ANOVA was used. When analyses involved multiple factors, a two-way ANOVA assessed the effects and their interactions, followed by Sidak’s or Dunnett’s multiple comparisons tests as indicated.

All in vitro experiments, including biochemical and enzymatic assays, were independently replicated at least once using separate batches of purified protein to confirm reproducibility. Each protein batch was derived from the same stable cell line but cultured on different occasions to constitute biological replicates. Results from biological replicates were consistent; thus, representative graphs are shown.

Statistical analyses, graphing, and enzyme kinetic evaluations were performed using GraphPad Prism v10.2. Figures were prepared and edited in Affinity Publisher v2.4.2. Some graphic schematics incorporated in the figures were sourced from Bioicons.com.

### Data availability

Coordinates and cryo-EM maps have been deposited in the PDB and EMDB, with the following accession numbers: apo-CLN8 (PDB: 9THK; and EMD: 55929), CLN8 + C18:1-CoA (PDB: 9TKA; and EMD: 56022), CLN8 + C22:6-CoA (PDB: 9TK9 and EMD: 56021). All data needed to evaluate the conclusions stated in the paper are presented in the paper and/or the supplementary materials. Requests for reagents or biological materials should be directed to the lead author.

## Acknowledgements

We thank Julian Rayner for advice; Dima Chirgadze, Lee Cooper, Giulia Paris, and Luay Joudeh from University of Cambridge Cryo-EM facility for cryo-EM support; Samuel Lord from CRUK Cambridge Institute Proteomics Core facility for proteomics analysis. R,S-GPG was synthesised by o2h Group and S,S-C18:1-LPG by WuXi AppTec. Gene synthesis was done by GenScript, and Sanger sequencing was done by Source Genomics. Manuscript preparation was assisted by ChatGPT 5 (OpenAI). This study was supported by grants from Medical Research Council (MC_UU_00028), Batten Disease Global Research Initiative (BDGRI), NCL Foundation (NCL-Stiftung), Italian Ministry of Health (RC 2025/2026), Telethon Foundation (GSA23C003), Wellcome Trust Investigator award (220257/Z/20/Z), the University of Oulu Graduate School, the Research Council of Finland (311934; 331436). K.P. has received a Springboard award from Wellcome Trust - Academy of Medical Sciences (AMS) (SBF0010/1078). M.M. is a former Telethon Career awardee.

## Author contributions

K.P. conceived and supervised the project. P.K.S., A.M.J., J.V.D.K., S.K.L., N.J. and K.P. designed and performed experiments. D.L., P.K.S., J.J.R., M.S.K. and E.R.S.K. prepared cryo-EM samples, analysed and interpreted data. A.M.J., B.J., C.S.Y., A.K. and K.P. performed mass- spectrometry analyses. S.D.V., F.M.S. and M.M. performed zebrafish studies. K.T. and K.P. performed mouse LPG experiment. J.M.N., M.H.S., M.A.P., J.U., J.M.W. and R.H. provided mouse tissues. J.U., J.M.W., A.K., R.H., F.M.S., M.M., M.P.M., E.R.S.K. and K.P. secured funding and provided overall supervision and advice. K.P. wrote the manuscript with input from all authors.

## Ethics declarations

The authors declare no competing interests.

## Description of Supplementary Tables

Supplementary Table 1. Cryo-EM data collection, refinement and validation statistics.

Supplementary Table 2. Details of antibodies used in this study.

Supplementary Table 3. Details of chemicals and reagents used in this study.

Supplementary Table 4. Details of plasmids used in this study.

Supplementary Table 5. MS/MS transitions (positive mode), cone voltages, and collision energies used in this study.

---

**Extended Data Fig. 1:**
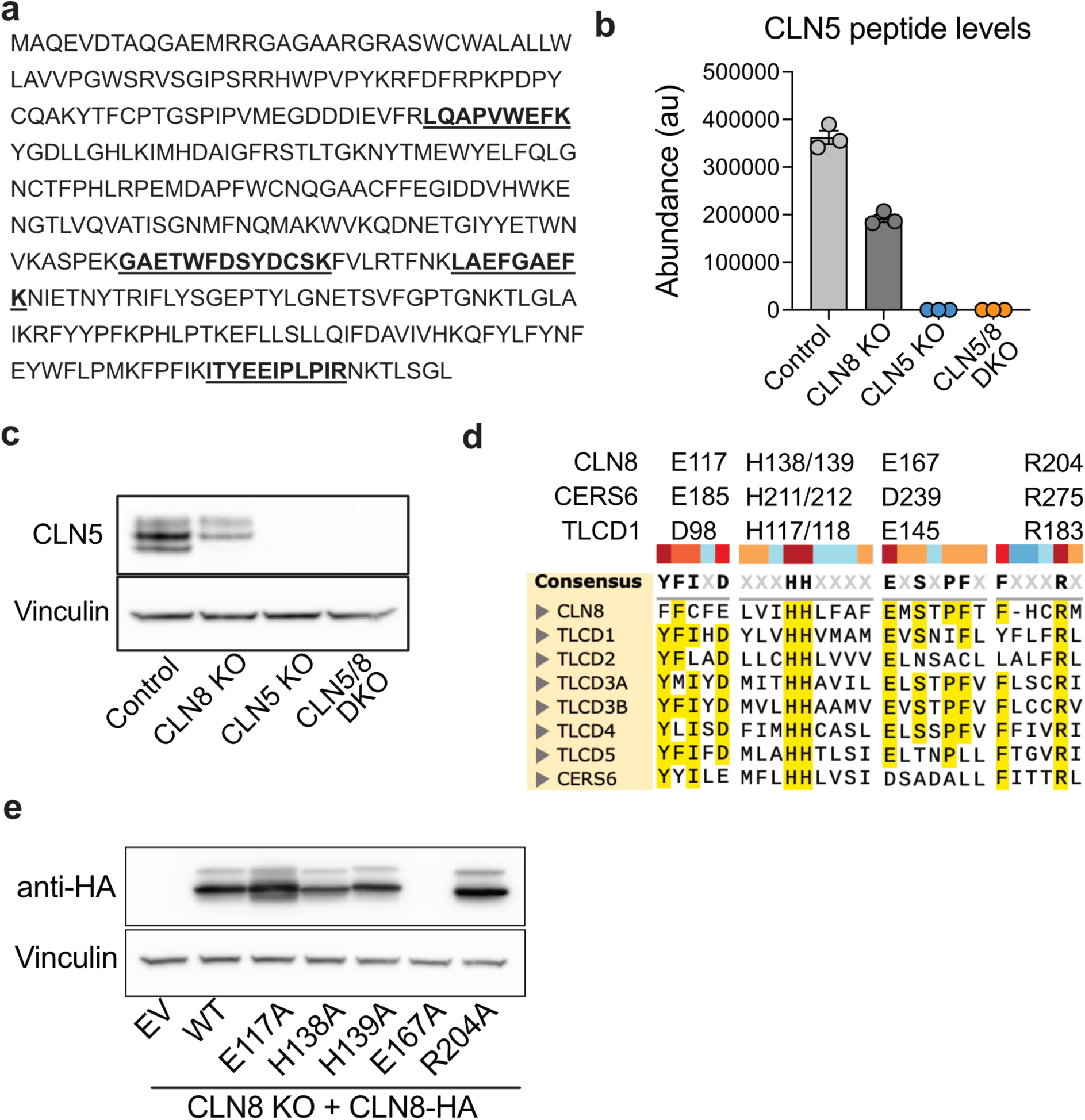
Validation of CLN5 and CLN8 HeLa cell models and overexpression constructs. a, CLN5 tryptic peptides used for protein quantification, highlighted within the CLN5 amino acid sequence. **b,** The abundance of CLN5 peptides in HeLa control, CLN8 KO, CLN5 KO, and CLN5/8 double KO (DKO) cell lines (n = 3). **c,** Immunoblot showing CLN5 protein levels and vinculin loading control in the same cell lines as in (**b**). **d,** Protein sequence alignment of selected TLCD family members, showing regions surrounding conserved residues only; residue numbers for CLN8, CERS6, and TLCD1 are indicated above. **e,** Expression of the indicated CLN8-HA mutant constructs in CLN8 KO cells.

**Extended Data Fig. 2:**
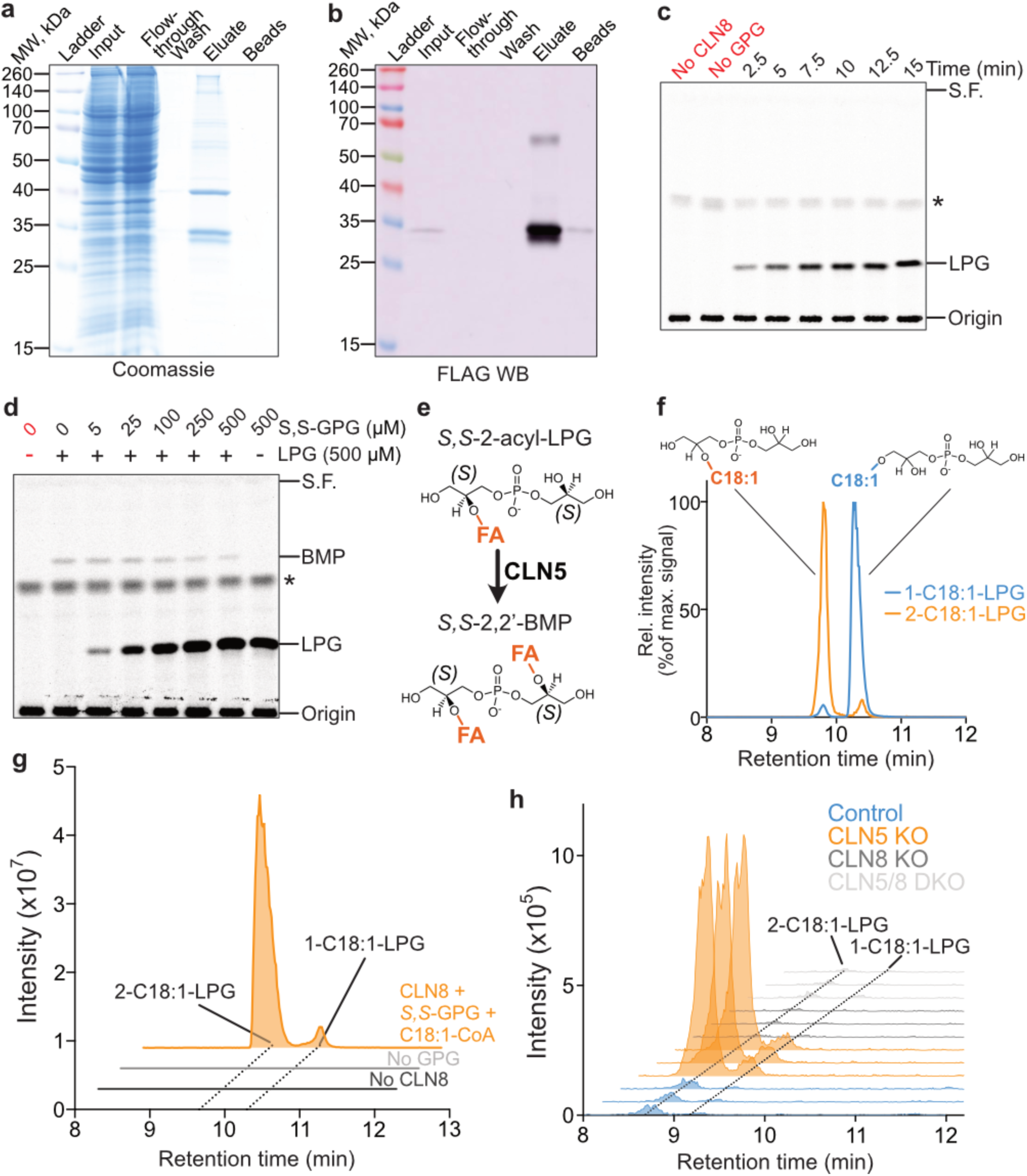
Purification and enzymatic characterization of CLN8 expressed in human cells. a, Coomassie-stained SDS–PAGE showing total protein in the indicated fractions collected during purification of FLAG–STREP-tagged CLN8. **b,** Western blot detecting the FLAG tag in the same fractions as in (**a**). The eluate, passed through size- exclusion spin columns to remove biotin, was used directly for in vitro assays. The blot is displayed as a chemiluminescence–colorimetric overlay. **c,** Phosphorimage of a TLC plate showing a time-course assay with purified CLN8 incubated with oleoyl-CoA containing a [1- ¹⁴C]-oleoyl-CoA tracer and *S,S*-GPG substrate. **d,** Phosphorimage of a TLC plate from a substrate-competition assay using purified CLN8 and [1-¹⁴C]-oleoyl-CoA, with *R,rac*-LPG fixed at 500 µM and *S,S*-GPG titrated at the indicated concentrations. In (**c,d**), migration positions of LPG and BMP were validated using chemical standards; the asterisk denotes a non- specific band from the [1-¹⁴C]-oleoyl-CoA preparation. **e,** Schematic illustrating the *sn-2* acyl positions of the LPG precursor and BMP product. **f,** LC-MS/MS chromatograms showing the migration of 1-C18:1-LPG and 2-C18:1-LPG standards. **g,** LC-MS/MS chromatogram of C18:1- LPG generated in vitro from purified CLN8, *S,S*-GPG, and C18:1-CoA, compared against positional isomer standards. **h,** LC–MS/MS chromatograms of C18:1-LPG detected in control, CLN8 KO, CLN5 KO, and CLN5/8 DKO HeLa cells (n = 3 per genotype), resolved against positional isomer standards. In vitro assay results are representative of two independent CLN8 purifications.

**Extended Data Fig. 3:**
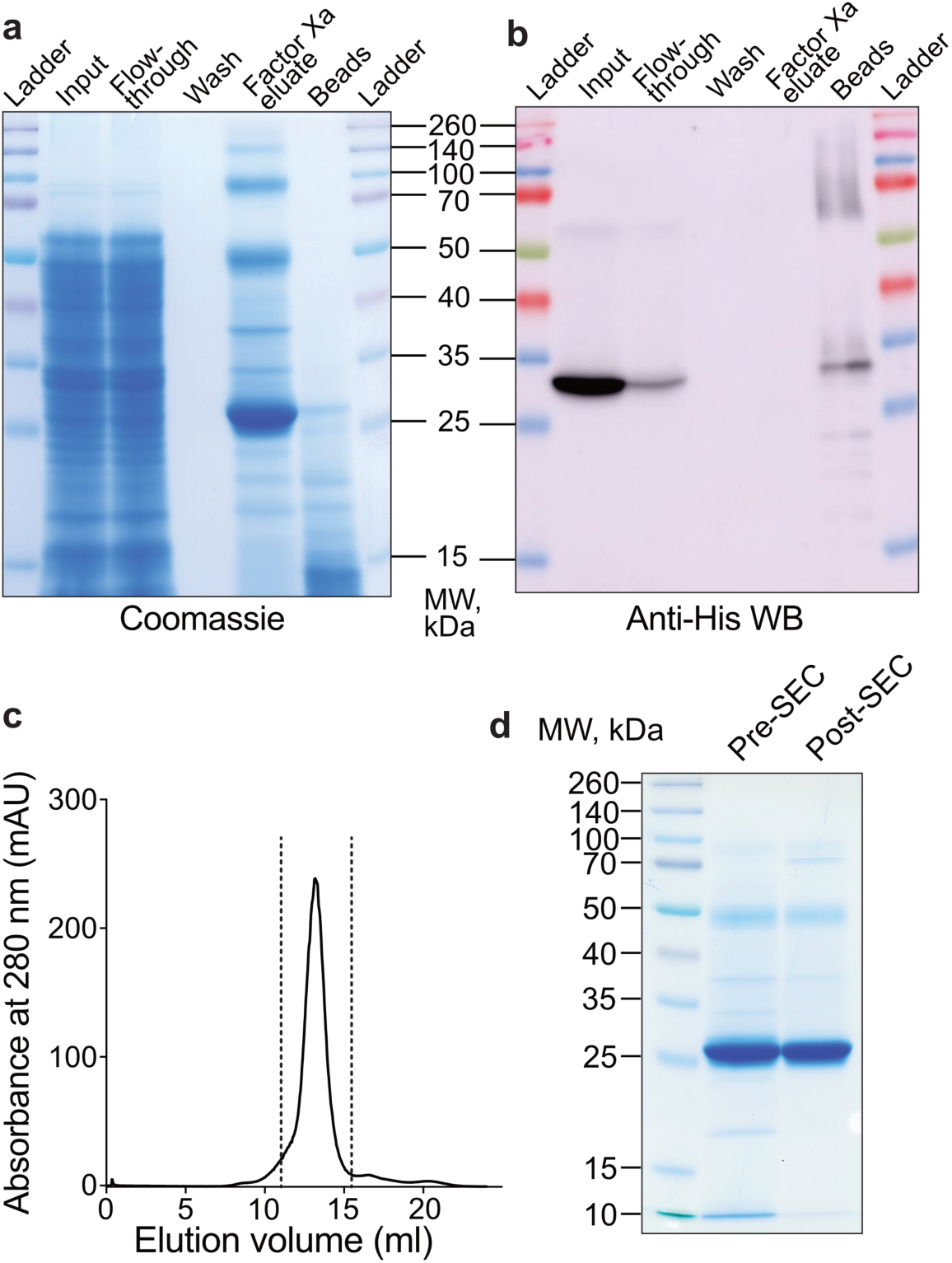
Purification of CLN8 from yeast for structural studies. **a**, Coomassie- stained SDS–PAGE showing total protein in the indicated fractions during purification of 6×His-tagged, codon-optimised CLN8, containing a Factor Xa protease cleavage site between the tag and coding sequence. **b**, Western blot detecting the 6×His tag in the same fractions as in (**a**). **c**, Size-exclusion chromatography (SEC) of the eluate from (**a**), showing a single homogeneous protein population. Dashed lines indicate the fractions collected for cryo-EM. **d**, Coomassie-stained SDS–PAGE comparing eluate before and after size-exclusion chromatography, with the post-SEC sample used for cryo-EM grid preparation.

**Extended Data Fig. 4:**
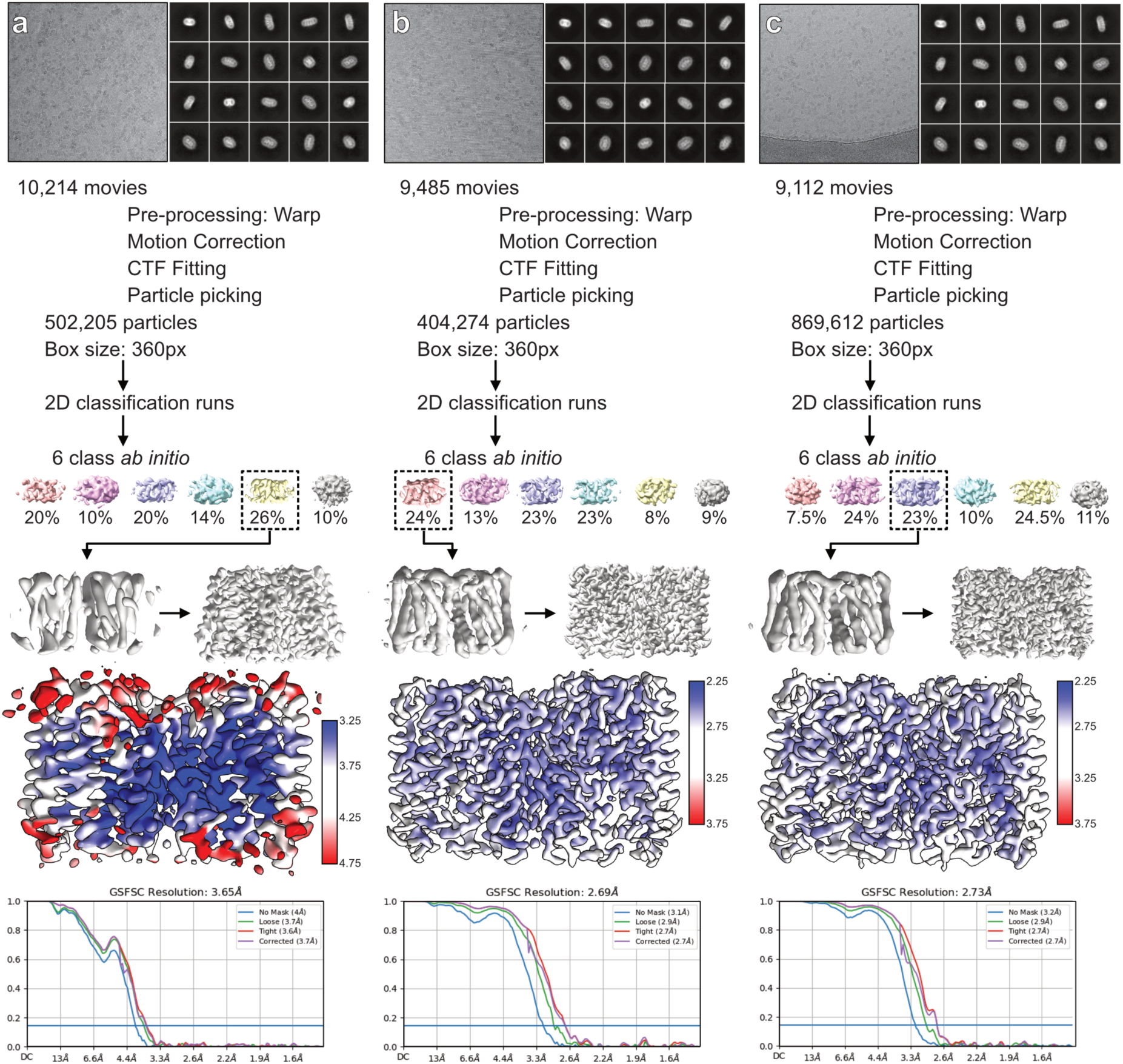
Cryo-EM data processing for CLN8 apo and substrate-bound structures. Representative micrographs, 2D class averages obtained prior to 3D classification, data-processing flow charts, final 3D reconstructions coloured by local resolution, and Fourier shell correlation (FSC) plots indicating overall map resolutions are shown for: **a**, apo CLN8 structure; **b**, CLN8 incubated with 100 µM C18:1-CoA for 1 h at room temperature; **c**, CLN8 incubated with 100 µM C22:6-CoA for 1 h at room temperature.

**Extended Data Fig. 5:**
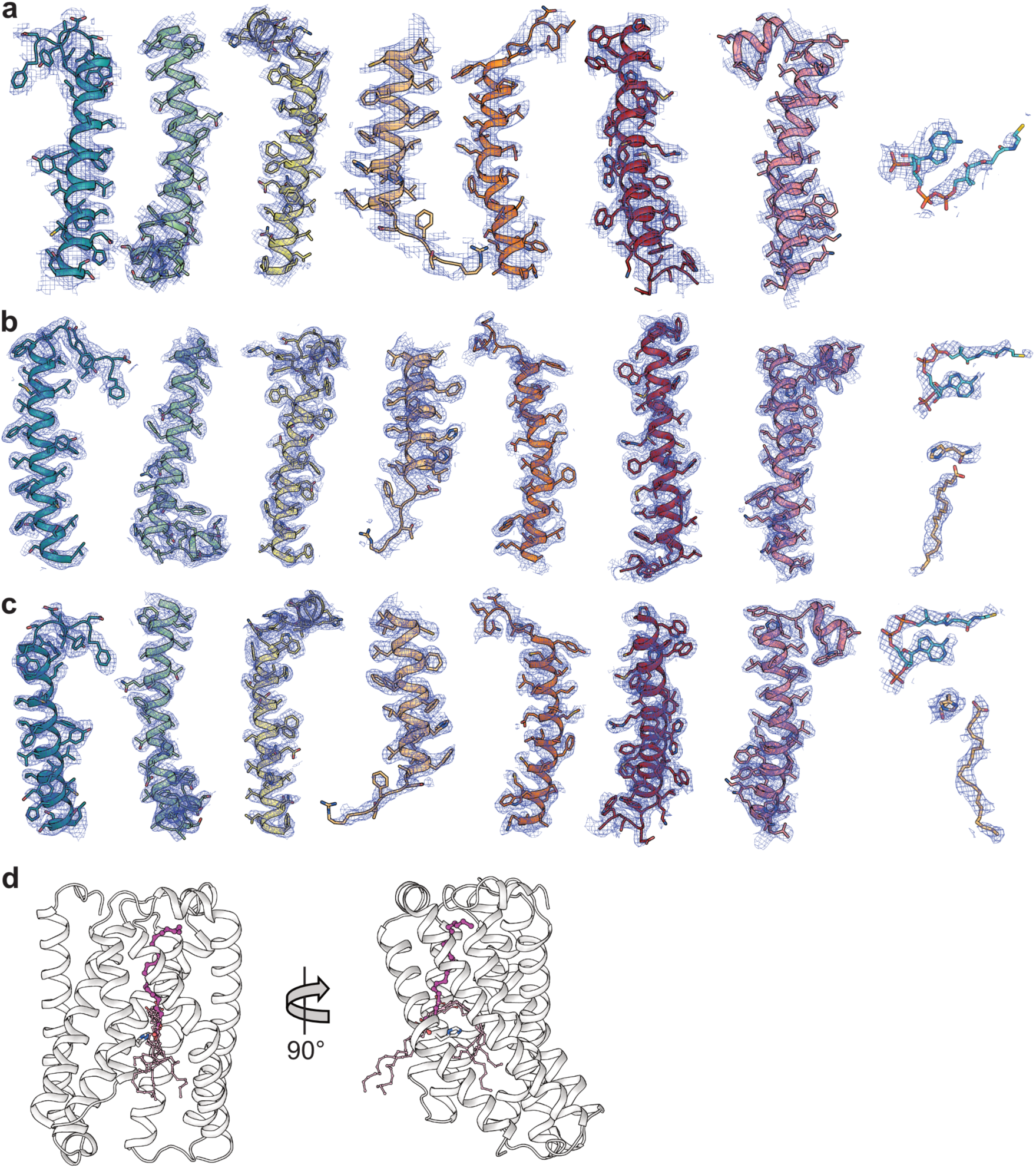
Cryo-EM map quality validation for CLN8 apo and substrate-bound structures. Representative cryo-EM densities (black mesh) for transmembrane helices and bound CoA, with side chains shown as sticks: **a**, apo CLN8; **b**, CLN8 solved with C18:1–CoA; **c**, CLN8 solved with C22:6–CoA. Maps are contoured in Chimera at 0.11 (0.56σ), 0.246 (1.68σ) and 0.22 (1.73σ), respectively, and are displayed within 2 Å of the depicted features. For (**b**,**c**), the positions of the fatty acyl chains adjacent to H138 are indicated. Note the absence of continuous density between the fatty acyl carboxyl groups and H138. **d**, Cartoon representation of the CLN8 structure solved with C22:6-CoA, showing the main observed density for C22:6 (red) and minor densities corresponding to alternative binding poses (yellow).

**Extended Data Fig. 6:**
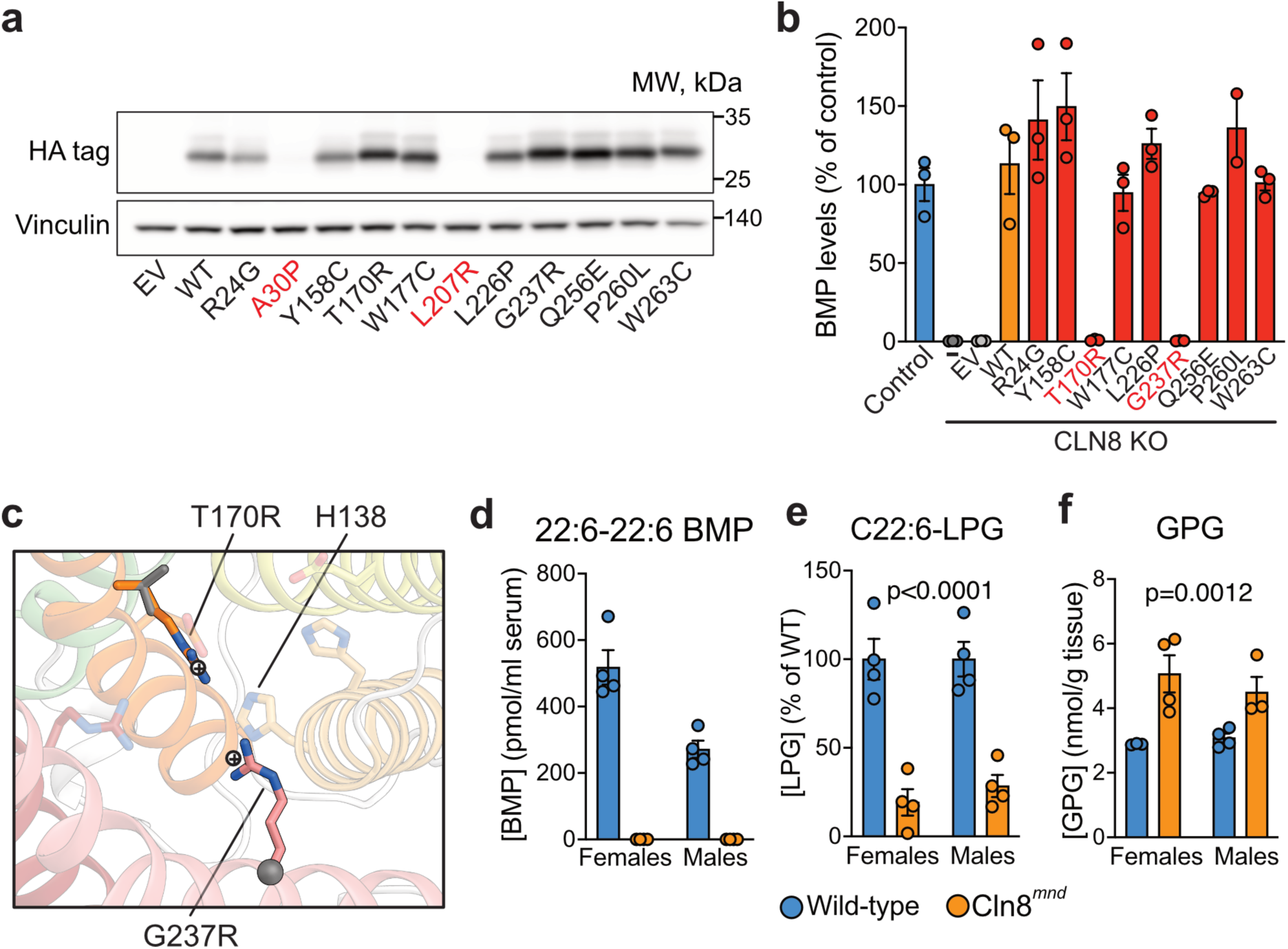
Expression and functional impact of CLN8 mutations. **a**, Western blot analysis of HA-tagged CLN8 constructs expressed in CLN8 KO HeLa cells, detected with anti- HA antibody, with vinculin as a loading control (48 h post-transfection). **b**, Total BMP levels measured in control or CLN8 KO HeLa cells transfected with the indicated CLN8-HA constructs (n = 3, 48 h). **c**, Structural overlay showing the positions of T170R and G237R mutations within the CLN8 active site. The side chains of the mutant arginines are positioned 2.7 Å and 4.3 Å from the catalytic residue H138, respectively. **d,** Levels of 22:6–22:6-BMP in the serum of Cln8*^mnd^* mice (n = 4 per sex). **e,** Levels of C22:6-LPG, and (**f**) GPG in the brains of wild-type and Cln8*^mnd^* mice (n = 3–4 per sex). In **(e,f)**, indicated *P* values reflect genotype effects in two-way ANOVA; the sex factor was not significant.

**Extended Data Fig. 7.**
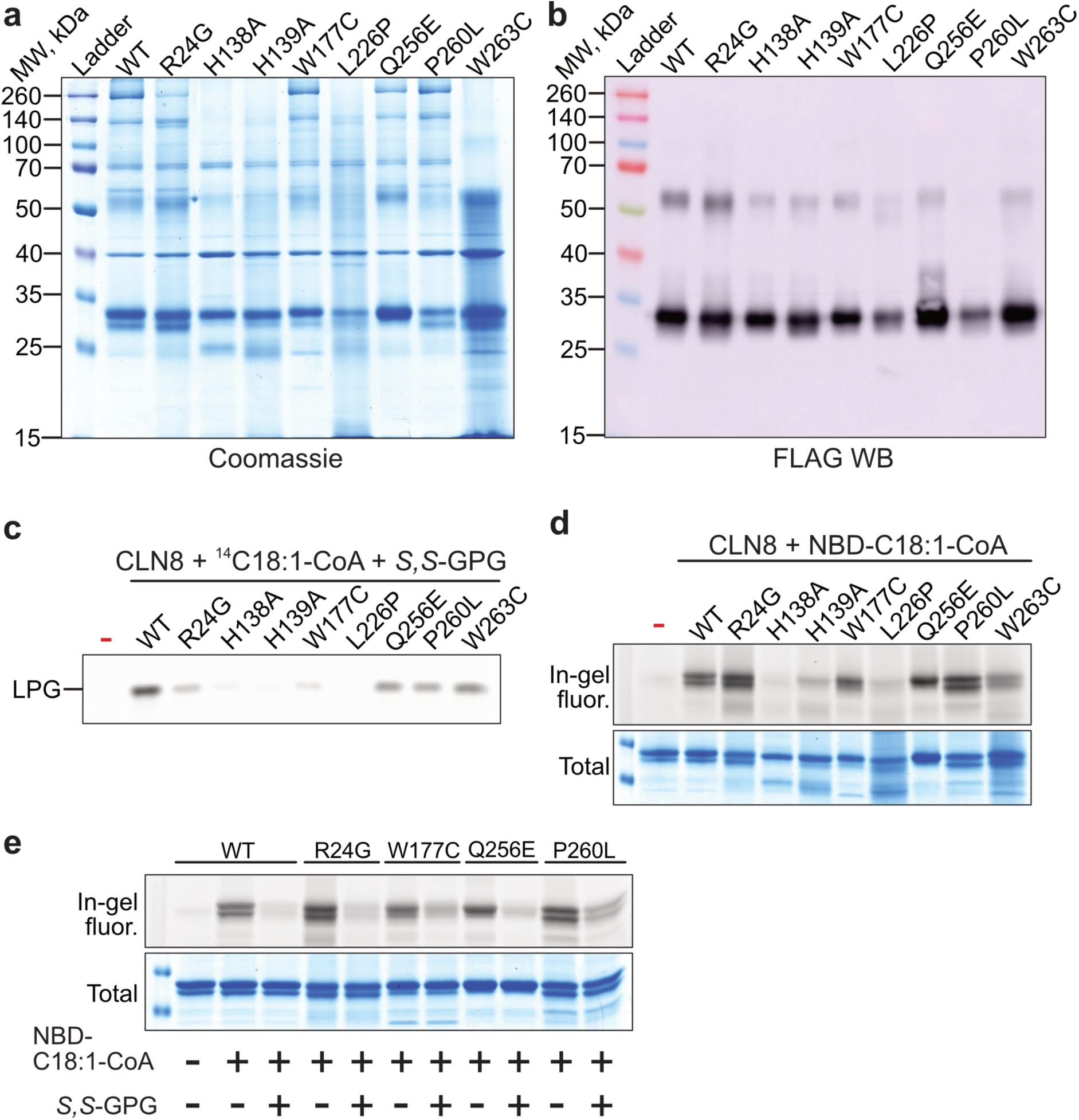
Purification and in vitro characterisation of CLN8 Batten disease mutant variants. **a**, Coomassie-stained SDS–PAGE showing total protein in eluate fractions from purification of FLAG–STREP–tagged CLN8 mutant variants. Eluates were passed through size-exclusion spin columns to remove biotin and used directly in in vitro assays. **b**, Western blot detection of FLAG in the same fractions as in (**a**). **c**, Phosphorimaging of a TLC plate showing enzymatic activity of purified CLN8 variants with oleoyl-CoA (with [1-¹⁴C]-oleoyl-CoA tracer) and *S,S*-GPG as substrates. **d**, In-gel NBD fluorescence (top) and Coomassie staining (bottom) of SDS–PAGE of purified CLN8 variants incubated with NBD–18:1–CoA alone. **e**, As in (**d**), but incubated with both NBD-18:1-CoA and *S,S*-GPG as indicated. Migration of LPG was validated using a chemical standard. All assays are representative of two independent protein purifications.

**Extended Data Fig. 7.**
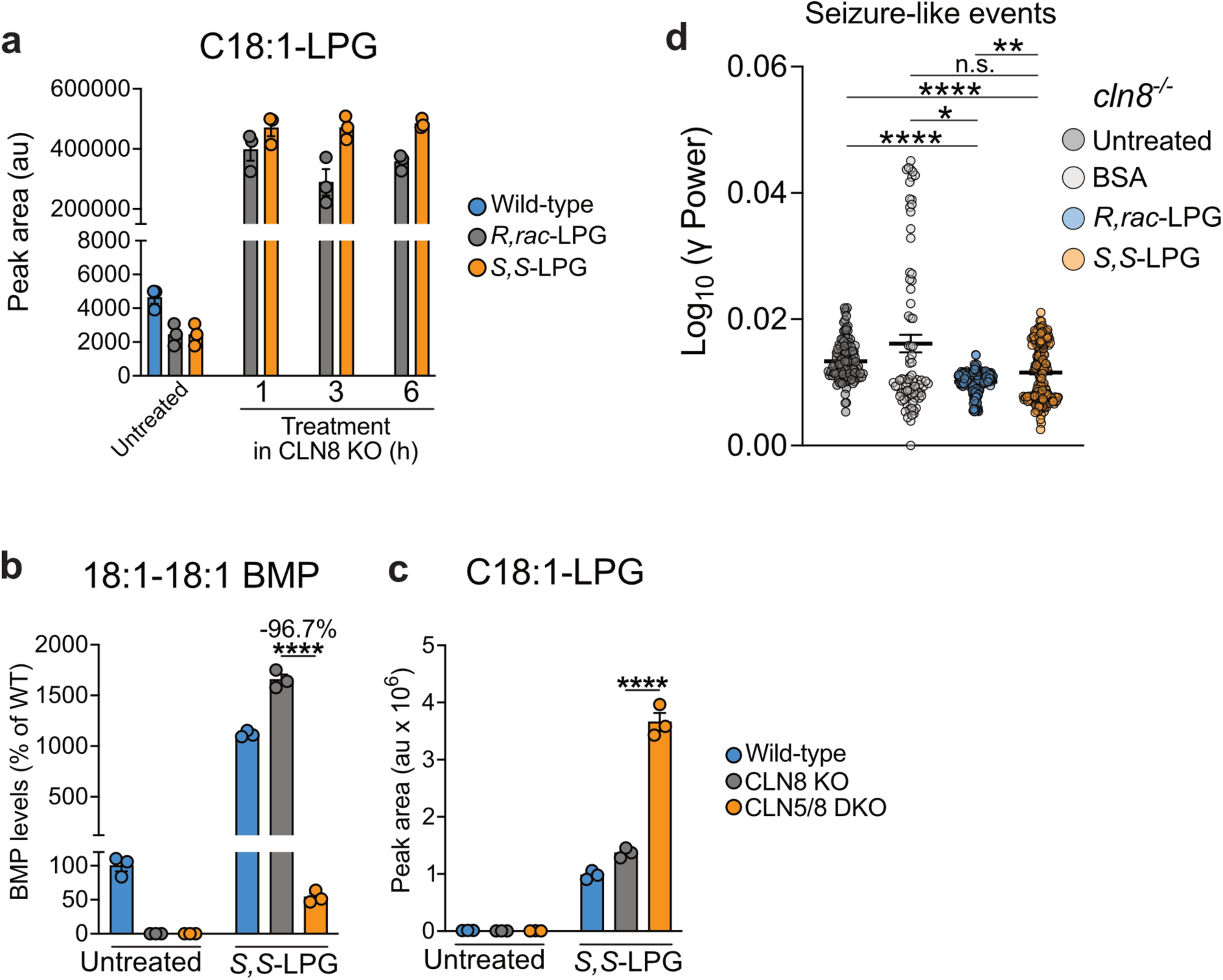
Properties of *S,S*-LPG-mediated rescue in CLN8-deficient models. a, Levels of C18:1-LPG species in wild-type and CLN8-knockout HeLa cells treated with 10 µM *R,rac*-LPG or *S,S*-LPG for the indicated times. Wild-type cells were untreated. **b,** Levels of 18:1-18:1 BMP and **(c)** C18:1-LPG species in wild-type, CLN8 KO, and CLN5/8 DKO HeLa cells untreated or treated with 10 µM *S,S*-LPG for 6 h. **d,** Power of seizure-like events detected by local field potential recordings at 120 hpf (n=8 larvae per group). In **(b,c)**, ****P < 0.0001 by two-way ANOVA with Sidak’s post-hoc test. In **(d)**, statistical analysis was performed using the Kruskal-Wallis test followed by Dunn’s post-hoc test (*p < 0.05; **p < 0.01; ***p < 0.001; ****p < 0.0001).

## References

1. Mole, S. E. & Cotman, S. L. Genetics of the neuronal ceroid lipofuscinoses (Batten disease). Biochim. Biophys. Acta BBA - Mol. Basis Dis. 1852, 2237–2241 (2015).

2. Sheokand, P. K. et al. TRAM-LAG1-CLN8 family proteins are acyltransferases regulating phospholipid composition. Sci. Adv. 11, eadr3723 (2025).

3. Medoh, U. N. et al. The Batten disease gene product CLN5 is the lysosomal bis(monoacylglycero)phosphate synthase. Science 381, 1182–1189 (2023).

4. Simonati, A. & Williams, R. E. Neuronal Ceroid Lipofuscinosis: The Multifaceted Approach to the Clinical Issues, an Overview. Front. Neurol. 13, (2022).

5. Medoh, U. N. & Abu-Remaileh, M. The Bis(monoacylglycero)-phosphate Hypothesis: From Lysosomal Function to Therapeutic Avenues. Annu. Rev. Biochem. 93, 447–469 (2024).

6. Brotherus, J., Renkonen, O., Herrmann, J. & Fischer, W. Novel stereoconfiguration in lyso- bis-phosphatidic acid of cultured BHK-cells. Chem. Phys. Lipids 13, 178–182 (1974).

7. Joutti, A., Brotherus, J., Renkonen, O., Laine, R. & Fischer, W. The stereochemical configuration of lysobisphosphatidic acid from rat liver, rabbit lung and pig lung. Biochim. Biophys. Acta 450, 206–209 (1976).

8. Somerharju, P., Brotherus, J., Kahma, K. & Renkonen, O. Stereoconfiguration of bisphosphatidic and semilysobisphosphatidic acids from cultured hamster fibroblasts (BHK cells). Biochim. Biophys. Acta 487, 154–162 (1977).

9. Somerharju, P. & Renkonen, O. Conversion of phosphatidylglycerol lipids to bis(monoacylglycero)phosphate in vivo. Biochim. Biophys. Acta 618, 407–419 (1980).

10. Frentzen-Bertrams, M. & Debuch, H. Production of bis(monoacylglycero)phosphate from phosphatidylglycerol in isolated liver lysosomes of chloroquine-pretreated rats. Hoppe. Seylers Z. Physiol. Chem. 362, 1229–1236 (1981).

11. Waite, M., Roddick, V., Thornburg, T., King, L. & Cochran, F. Conversion of phosphatidylglycerol to lyso(bis)phosphatidic acid by alveolar macrophages. FASEB J. Off. Publ. Fed. Am. Soc. Exp. Biol. 1, 318–325 (1987).

12. Waite, M., King, L., Thornburg, T., Osthoff, G. & Thuren, T. Y. Metabolism of phosphatidylglycerol and bis(monoacylglycero)-phosphate in macrophage subcellular fractions. J. Biol. Chem. 265, 21720–21726 (1990).

13. Thornburg, T., Miller, C., Thuren, T., King, L. & Waite, M. Glycerol reorientation during the conversion of phosphatidylglycerol to bis(monoacylglycerol)phosphate in macrophage- like RAW 264.7 cells. J. Biol. Chem. 266, 6834–6840 (1991).

14. Bulfon, D. et al. Functionally overlapping intra- and extralysosomal pathways promote bis(monoacylglycero)phosphate synthesis in mammalian cells. Nat. Commun. 15, 9937 (2024).

15. Głąb, B. et al. Cloning of Glycerophosphocholine Acyltransferase (GPCAT) from Fungi and Plants: A NOVEL ENZYME IN PHOSPHATIDYLCHOLINE SYNTHESIS. J. Biol. Chem. 291, 25066–25076 (2016).

16. Stålberg, K., Neal, A. C., Ronne, H. & Ståhl, U. Identification of a novel GPCAT activity and a new pathway for phosphatidylcholine biosynthesis in *S. cerevisiae*. J. Lipid Res. 49, 1794–1806 (2008).

17. Hejazian, S. M., Pirmoradi, S., Zununi Vahed, S., Kumar Roy, R. & Hosseiniyan Khatibi, S. M. An update on Glycerophosphodiester Phosphodiesterases; From Bacteria to Human. Protein J. 43, 187–199 (2024).

18. Nyame, K. et al. PLA2G15 is a BMP hydrolase and its targeting ameliorates lysosomal disease. Nature 642, 474–483 (2025).

19. Pascoa, T. C. et al. Structural basis of the mechanism and inhibition of a human ceramide synthase. Nat. Struct. Mol. Biol. (2024) doi:10.1038/s41594-024-01414-3.

20. Lonka, L., Kyttälä, A., Ranta, S., Jalanko, A. & Lehesjoki, A.-E. The neuronal ceroid lipofuscinosis CLN8 membrane protein is a resident of the endoplasmic reticulum. Hum. Mol. Genet. 9, 1691–1697 (2000).

21. Katata, Y. et al. Novel missense mutation in CLN8 in late infantile neuronal ceroid lipofuscinosis: The first report of a CLN8 mutation in Japan. Brain Dev. 38, 341–345 (2016).

22. Cannelli, N. et al. Novel mutations in CLN8 in Italian variant late infantile neuronal ceroid lipofuscinosis: Another genetic hit in the Mediterranean. Neurogenetics 7, 111–117 (2006).

23. Siintola, E., Lehesjoki, A.-E. & Mole, S. E. Molecular genetics of the NCLs -- status and perspectives. Biochim. Biophys. Acta 1762, 857–864 (2006).

24. Sahin, Y. et al. Exome sequencing identifies a novel homozygous CLN8 mutation in a Turkish family with Northern epilepsy. Acta Neurol. Belg. 117, 159–167 (2017).

25. Ranta, S. et al. Variant late infantile neuronal ceroid lipofuscinosis in a subset of Turkish patients is allelic to Northern epilepsy. Hum. Mutat. 23, 300–305 (2004).

26. Badura-Stronka, M. et al. CLN8 Mutations Presenting with a Phenotypic Continuum of Neuronal Ceroid Lipofuscinosis-Literature Review and Case Report. Genes 12, 956 (2021).

27. Mahajnah, M. & Zelnik, N. Phenotypic heterogeneity in consanguineous patients with a common CLN8 mutation. Pediatr. Neurol. 47, 303–305 (2012).

28. Kolesnikova, M., et al. Phenotypic Variability of Retinal Disease Among a Cohort of Patients With Variants in the CLN Genes. Invest. Ophthalmol. Vis. Sci. 64, 23 (2023).

29. Ranta, S. & Lehesjoki, A.-E. Northern epilepsy, a new member of the NCL family. Neurol. Sci. 21, S43–S47 (2000).

30. Reinhardt, K. et al. Novel CLN8 mutations confirm the clinical and ethnic diversity of late infantile neuronal ceroid lipofuscinosis. Clin. Genet. 77, 79–85 (2010).

31. Kousi, M., Lehesjoki, A.-E. & Mole, S. E. Update of the mutation spectrum and clinical correlations of over 360 mutations in eight genes that underlie the neuronal ceroid lipofuscinoses. Hum. Mutat. 33, 42–63 (2012).

32. Ranta, S. et al. Variant late infantile neuronal ceroid lipofuscinosis in a subset of Turkish patients is allelic to Northern epilepsy. Hum. Mutat. 23, 300–305 (2004).

33. Messer, A., Strominger, N. L. & Mazurkiewicz, J. E. Histopathology of the late-onset motor neuron degeneration (Mnd) mutant in the mouse. J. Neurogenet. 4, 201–213 (1987).

34. Messer, A., Plummer, J., Maskin, P., Coffin, J. M. & Frankel, W. N. Mapping of the motor neuron degeneration (Mnd) gene, a mouse model of amyotrophic lateral sclerosis (ALS). Genomics 13, 797–802 (1992).

35. Ranta, S. et al. The neuronal ceroid lipofuscinoses in human EPMR and mnd mutant mice are associated with mutations in CLN8. Nat. Genet. 23, 233–236 (1999).

36. Müller-Niva, J. et al. Novel Cln8 p.R24G mouse line replicates major clinical features of Northern epilepsy. 2025.12.02.690107 Preprint at 10.64898/2025.12.02.690107 (2025).

37. Marchese, M. et al. Targeting autophagy impairment improves the phenotype of a novel CLN8 zebrafish model. Neurobiol. Dis. 197, 106536 (2024).

38. Breithofer, J., et al. CLN8 enables a non-canonical phospholipid synthesis pathway. bioRxiv (2025).

39. Laqtom, N. N. et al. CLN3 is required for the clearance of glycerophosphodiesters from lysosomes. Nature 609, 1005–1011 (2022).

40. Hobert, J. A. & Dawson, G. A novel role of the Batten disease gene CLN3: Association with BMP synthesis. Biochem. Biophys. Res. Commun. 358, 111–116 (2007).

41. Westerfield, M. The Zebrafish Book: A Guide for the Laboratory Use of Zebrafish (Danio Rerio). (University of Oregon Press, 2000).

42. Della Vecchia, S., et al. Trehalose Treatment in Zebrafish Model of Lafora Disease. Int. J. Mol. Sci. 23, 6874 (2022).

43. King, M. S. & Kunji, E. R. S. Expression and Purification of Membrane Proteins in Saccharomyces cerevisiae. Methods Mol. Biol. Clifton NJ 2127, 47–61 (2020).

44. Tegunov, D. & Cramer, P. Real-time cryo-electron microscopy data preprocessing with Warp. Nat. Methods 16, 1146–1152 (2019).

45. Varadi, M. et al. AlphaFold Protein Structure Database: massively expanding the structural coverage of protein-sequence space with high-accuracy models. Nucleic Acids Res. 50, D439–D444 (2022).

46. Jumper, J. et al. Highly accurate protein structure prediction with AlphaFold. Nature 596, 583–589 (2021).

47. Pettersen, E. F. et al. UCSF Chimera--a visualization system for exploratory research and analysis. J. Comput. Chem. 25, 1605–1612 (2004).

48. Croll, T. I. ISOLDE: a physically realistic environment for model building into low- resolution electron-density maps. Acta Crystallogr. Sect. Struct. Biol. 74, 519–530 (2018).

49. Meng, E. C. et al. UCSF ChimeraX: Tools for structure building and analysis. Protein Sci. Publ. Protein Soc. 32, e4792 (2023).

50. Emsley, P., Lohkamp, B., Scott, W. G. & Cowtan, K. Features and development of Coot. Acta Crystallogr. D Biol. Crystallogr. 66, 486–501 (2010).

51. Sweeney, A., Mulvaney, T., Maiorca, M. & Topf, M. ChemEM: Flexible Docking of Small Molecules in Cryo-EM Structures. J. Med. Chem. 67, 199–212 (2024).

52. Davis, I. W. et al. MolProbity: all-atom contacts and structure validation for proteins and nucleic acids. Nucleic Acids Res. 35, W375–W383 (2007).

53. Yamashita, K., Palmer, C. M., Burnley, T. & Murshudov, G. N. Cryo-EM single-particle structure refinement and map calculation using Servalcat. Acta Crystallogr. Sect. Struct. Biol. 77, 1282–1291 (2021).

54. Tsuji, Y. Transmembrane protein western blotting: Impact of sample preparation on detection of SLC11A2 (DMT1) and SLC40A1 (ferroportin). PLoS ONE 15, e0235563 (2020).

55. Kawana, H. et al. An accurate and versatile method for determining the acyl group- introducing position of lysophospholipid acyltransferases. Biochim. Biophys. Acta BBA - Mol. Cell Biol. Lipids 1864, 1053–1060 (2019).

56. Löfgren, L., Forsberg, G.-B. & Ståhlman, M. The BUME method: a new rapid and simple chloroform-free method for total lipid extraction of animal tissue. Sci. Rep. 6, 27688 (2016).

57. Okudaira, M. et al. Separation and quantification of 2-acyl-1-lysophospholipids and 1- acyl-2-lysophospholipids in biological samples by LC-MS/MS. J. Lipid Res. 55, 2178–2192 (2014).

